# Targeting diverse operational regimes in recurrent spiking networks

**DOI:** 10.1101/2022.04.22.489005

**Authors:** Pierre Ekelmans, Nataliya Kraynyukova, Tatjana Tchumatchenko

**Affiliations:** Max Planck Institute for Brain Research Theory of neural dynamics group; Frankfurt Institute for Advanced Studies; University of Bonn Medical Center Institute of experimental epileptology and cognition research; University of Mainz Medical Center Institute of physiological chemistry

## Abstract

Neural computations emerge from recurrent neural circuits that comprise hundreds to a few thousand neurons. Continuous progress in connectomics, electrophysiology, and calcium imaging require tractable spiking network models that can consistently incorporate new information about the network structure and reproduce the recorded neural activity features. However, it is challenging to predict which spiking network connectivity configurations and neural properties can generate fundamental operational states and specific experimentally reported nonlinear cortical computations. Theoretical descriptions for the computational state of cortical spiking circuits are diverse, including the balanced state where excitatory and inhibitory inputs balance almost perfectly or the inhibition stabilized state (ISN) where the excitatory part of the circuit is unstable. It remains an open question whether these states can co-exist with experimentally reported nonlinear computations and whether they can be recovered in biologically realistic implementations of spiking networks. Here, we show how to identify spiking network connectivity patterns underlying diverse nonlinear computations such as XOR, bistability, inhibitory stabilization, supersaturation, and persistent activity. We established a mapping between the stabilized supralinear network (SSN) and spiking activity which allowed us to pinpoint the location in parameter space where these activity regimes occur. Notably, we found that biologically-sized spiking networks can have irregular asynchronous activity that does not require strong excitation-inhibition balance or large feedforward input and we showed that the dynamic firing rate trajectories in spiking networks can be precisely targeted without error-driven training algorithms.

## I. INTRODUCTION

Layered or columnar neuronal structures consisting of hundreds to thousands of neurons constitute local computational blocks in the mammalian cortex. Each computational block has its particular size and connectivity rules, which determine its dynamics and computational repertoire. Therefore, understanding the computational regimes of recurrent networks with different sizes ranging from hundreds to millions of neurons and permitting diverse connectivity patterns is essential to explain the emergence of cognitive functions and behavior. It is currently a challenge to quantitatively predict the activity of medium-sized spiking neural networks. The most powerful mathematical theories now operate at two opposite scales: they either model a small number of neurons that generate a particular activity pattern of interest [1], or they operate in the limit of large or infinitely large networks [2]. Few theories can make mathematically tractable, quantitative, and experimentally relevant predictions for the diverse, intermediate sizes of spiking networks reported for local cortical circuits [3–5].

Here, we study the activity regimes of spiking networks whose sizes range from a few hundred to thousands of neurons. Many parameters describing spiking neurons and their intracortical connections have recently been measured across cortical cell types [6], and detailed numerical network simulations have been put forward [7]. However, it is challenging to interpret spiking network simulations because network dynamics depend strongly on multidimensional parameter settings, while experimentally reported parameters can vary within broad ranges [8–12]. On the opposite side of the complexity spectrum are population rate networks [13–15] which describe the average activity of neurons in each population and can relate a specific activity regime to a connectivity configuration. However, it is still unclear how to translate computational results obtained with rate models to biologically plausible spiking networks.

One of the most popular models for large or infinitely large networks is the balanced state framework [16]. Its biological correlate is the experimentally reported strong balance between excitatory and inhibitory synaptic currents [17] and asynchronous irregular spiking activity [18]. However, the computational hallmarks of the balanced network limit, including response linearity and strong feedforward connections, are not consistent with the diversity of experimentally reported non-linear responses across cortical areas [19] and reports of weak feedforward inputs [20]. Therefore, experimental observations deviating from the balanced state predictions have been often addressed by keeping the core excitatory/inhibitory (E/I) balance framework while adding specific synaptic plasticity rules [21], more complex synaptic strength distributions [22], or a semi-balance condition which allows for excess inhibition [23]. Furthermore, the existence and stability of the balanced state limit come with strict conditions on the feedforward and recurrent connection strengths (see [16, 24], Appendix), which exclude many network connectivity configurations. Finally, the balanced limit is based on the assumption that networks are infinitely large and ignores finite-size effects.

Firing rate models of neural sub-populations with a nonlinear activation function [13, 25, 26] are an alternative to the balanced state model. These models often use either a threshold linear function [25] or an S-shaped sigmoidal function [13, 27] to describe the activation function (F-I curve) of neurons. The advantage of the threshold linear network model is that it can be solved analytically; however, it fails to reproduce non-linear responses often observed in experiments. On the other hand, network models with the sigmoidal transfer functions support nonlinear responses; however, systematic mathematical analysis for the activity states of these networks is currently in short supply. The stabilized supralinear network model (SSN) [26] uses a supra-linear power law as a transfer function. The advantage of the SSN model is that its activity states can be characterized analytically [15, 28] and it can reproduce a variety of nonlinear cortical responses in the realistic range of firing rates observed *in vivo* [26].

Could the SSN model predict and quantify the activity regimes in medium-sized spiking networks for all connectivity configurations? Here, we argue that the SSN model can be used to predict diverse nonlinear responses such as supersaturation, bistable activity, and inhibition stabilized regimes in spiking networks. We propose a mapping between the parameter space of the SSN model and the more complex parameter space of the leaky-integrate-and-fire (LIF) network of spiking neurons. The advantage of this approach is that it results in a mathematically tractable model which can be manipulated analytically. Thanks to this, we can invert the model equations and design the network’s input to target a desired activity trajectory or operational regime of interest via closed-form equations without network training. This approach can be used to generate specific nonlinear functions (eg: XOR gate).

In relation to the balanced state, we show that for biologically realistic connectivity configurations, even very large networks (size *N* ≈ 10^5^) can deviate substantially from the balanced state predictions despite high E-I correlations. Likewise, networks lacking precise E-I input balance can produce asynchronous irregular firing patterns resembling balanced network activity. Overall, we find that medium-sized spiking networks can have a complex, nonlinear behavior that is accurately described with the SSN framework and can be far from the predictions of the balanced state limit.

Examining the possible activity regimes of networks with biologically plausible synaptic strengths, we could delineate different computational regimes such as supersaturation and bistability. We found that only a specific range of connectivity and input choices allow the networks to be in the inhibition stabilized state [25]. The inhibition stabilized state has been suggested as an operational state of the cortex [25, 29] and is linked to a decrease of the inhibitory firing rates following the increase in inhibitory input. Here, we delineate the parameter regimes in the medium-sized networks where inhibitory stabilization and a set of other experimentally reported nonlinear computations co-exist and parameter regions where they are mutually exclusive.

## II. RESULTS

Our goal is to understand how neural circuits comprising a few thousand neurons organize their spiking activity. We want to predict whether specific nonlinear computations can occur in these networks and pinpoint their location in the multidimensional parameter space spanned by recurrent connectivity and input weights. We choose the size of the networks to be 4000 neurons, which is biologically plausible for local cortical circuits (Appendix). Similarly, we restrict our analysis to the *in vivo* reported range of activity of 0-10 Hz [30–37]. We choose the strength and probability of synaptic connections to be within the same order of magnitude as the values reported by the database of the Allen Institute for the visual cortex area V1 in mice [6] (Appendix). To model cortical activity, we use the leaky-integrate-and-fire (LIF) model (see Methods), which represents an accurate description of cortical neurons both *in vivo* and *in vitro* [38]. To predict spiking network activity regimes, we map the mean activity of spiking networks to a rate-based 2D SSN model.

### A. Approximating spiking network activity with the SSN model

Our starting point is the observation that a power-law function can accurately describe the F-I curve of a single LIF neuron across different membrane time constants *τ* and input noise values *σ* (see Fig. **1B,C**). The power-law approximation is consistent with the Ricciardi exact solution of the LIF neuron Eq. **9** [39], which we denote by Φ (Fig. **1B**). Throughout this study, we are interested in the firing rate range from zero to approximately 10 Hz which has been reported *in vivo* across many brain areas [30–37]. To this end, we fit the low firing rate regime (0-10 Hz) of the F-I curve of a LIF neuron given by Φ using the threshold power-law function of the form

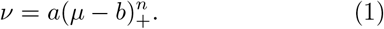

**FIG. 1.**
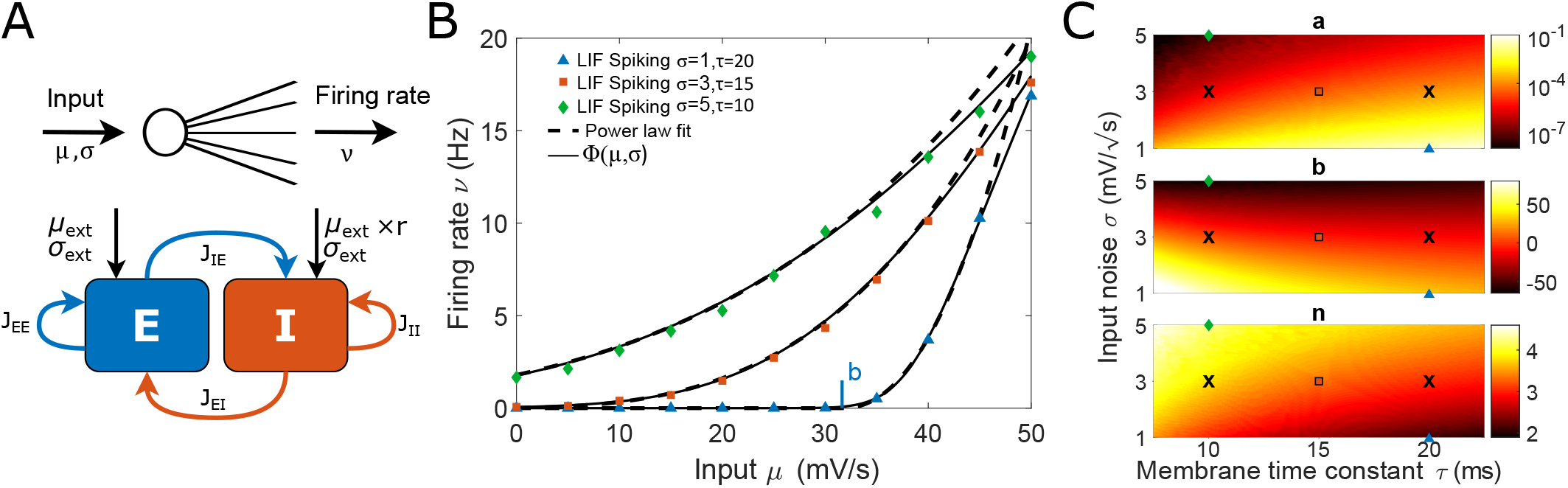
Spiking neurons can be quantitatively described by a supralinear power law for low activity. (A, top) Schematic representation of the F-I transfer function of a neuron. (A, bottom) Architecture of the recurrent Excitatory-Inhibitory network. (B) Neuronal firing rate as a function of input for different input noise *σ* and membrane time constant *τ*. The firing rate response of simulated LIF neurons (colored triangles) is in line with the Ricciardi solution Φ (solid line, Eq. **9**). The power-law approximation (Eq. **1**) accurately aligns with the LIF simulation and Ricciardi solution for low firing rates. Note that the power-law fit is only applied in the range of *ν <* 10 Hz, and diverges beyond this range. The vertical mark denotes the power-law parameter *b* for one of the curves (*b* is negative for the other two curves). (C) The power-law parameters depend on input noise *σ* and the membrane time constant *τ*. The two crosses indicate the parameter regimes we use for the excitatory (*τ*_*E*_ = 20 ms) and inhibitory (*τ*_*I*_ = 10 ms) neurons in recurrent networks. Other symbols indicate the parameters *τ* and *σ* used in B. The fit is obtained with the least squares method. Power-law parameters are listed in Table **III**.

Where *ν* is the firing rate, *µ* is the input to the neuron and (*x*)_+_ = max {*x*, 0}. The constants *a, b*, and *n* characterize the power-law approximation, with a scaling pre-factor *a*, an input threshold *b* upon which the neuron starts firing, and an exponent *n*. The power-law exponent *n* in our approximation varies between 2 and 4, which is consistent with the biologically reported range [35, 40].

We connect the individual LIF neurons into a recurrent network of excitatory (E) and inhibitory (I) neurons (Fig. **1A**). We assume that E and I neurons differ in their membrane time constants (*τ*_*E*_=20 ms, *τ*_*I*_ =10 ms, black crosses in Fig. **1C**) consistently with experimental reports [6]. We note that the input to a neuron in a recurrent network, which is a superposition of postsynaptic potentials (PSPs), is equivalent to an Ornstein Uhlenbeck process or white noise if the number of incoming PSPs is sufficiently large and the activity is irregular [41, 42].

To describe the activity of the E and I populations, we use the power-law approximation of the single-neuron transfer function (Eq. **1**). In the mean field approximation, the average firing rate of each population is given by a system of equations equivalent to the SSN [26]

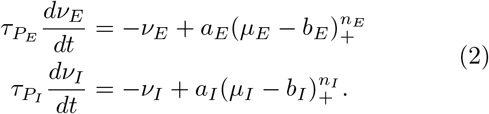

Where *ν*_*X*_, *X* ∈ {*E, I*} are the firing rates of the two populations, the parameters *a*_*X*_, *b*_*X*_, and *n*_*X*_ are given by the F-I curve fit of E and I neurons (black crosses in Fig. **1C**), and *µ*_*X*_ represent the sum of recurrent and feedforward inputs to each population. The population time constants 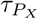 characterize how fast the firing rate of each population evolves. *In vivo* cortical networks have been shown to respond to sudden stimulation with a transient (or *onset*) response that has a time scale of approximately 20 ms [43]. Furthermore, *in vivo* recordings of multiple cortical areas have reported autocorrelation timescales of the order of hundreds ms [44–46] suggesting that slow timescales that are larger than the neuronal or synaptic variables are relevant for the analysis of biological circuits. Therefore, in the following we will consider firing rate dynamics that evolve on the time scales of a few tens to a hundred milliseconds which is slower than the intrinsic times scales of the network. Based on these time scales, the firing rate dynamics of the recurrent network can be understood using the equilibrium state and the external drive of a network. The equilibrium states of the SSN equation are described by

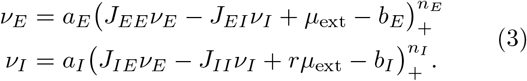

Here *r* is the ratio of the external inputs to the I and E populations *r* = *µ*_extI_*/µ*_extE_, which allows for the simplified notation: *µ*_ext_ = *µ*_extE_ and *rµ*_ext_ = *µ*_extI_. The population-wise connection strengths *J*_*XY*_ characterize the recurrent connections from population *Y* to population *X*, whereby *X, Y* ∈ {*E, I*}.

Previous work identified the constraints on connectivity configurations in the SSN model that underlie such nonlinear activity responses as supersaturation [15], the paradoxical effect [47, 48], bistability, and persistent activity [28]. We show that the parameters of LIF spiking networks can be mapped to the SSN such that the same activity types emerge in the spiking network, according to the observations made with the SSN. In the following sections, we discuss each activity type and its corresponding connectivity regime in the SSN, as well as in LIF spiking networks.

### B. Supersaturation - Firing rates can decline for growing input

Firing rates of neurons *in vivo* can show a range of non-linear behaviors as a function of stimulus strength [49]. In particular, the activity level of sensory neurons may decrease after stimulus onset, and a substantial number of pyramidal V1 neurons in mice show reduced firing in response to enhanced stimulus contrast [50]. At the same time, the average activity of thalamic neurons in mice - primarily targeting V1 neurons - is an increasing function of the stimulus contrast [51]. Therefore, it appears that E neurons can be suppressed despite the increase in external input. This phenomenon is generally referred to as supersaturation [15].

First, we studied E firing response to growing inputs and aimed to delineate parameter regimes where a decreasing population response can be observed. We found that supersaturation 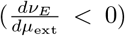 can be observed in a large class of connectivity and input weights configurations within the spiking networks that can be predicted by the inequality derived for the SSN model in [15, 52]

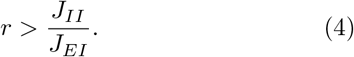

Interestingly, only three network parameters determine the SSN network’s ability to be in a supersaturating activity regime (Eq. **4**). For a network to be supersaturating, the ratio of external inputs *r* has to exceed the ratio of recurrent inhibition *J*_*II*_*/J*_*EI*_. As a result, the remaining two connectivity parameters (the recurrent excitation *J*_*IE*_ and *J*_*EE*_) cannot control the existence of supersaturating activity in the SSN. The exact point at which a network satisfying Eq. **4** becomes supersaturating does, however, depend on all network parameters as it occurs when the inhibitory firing rate exceeds a specific threshold value (Appendix, Eq. **B5**).

To understand if the connectivity condition derived in the SSN model (Eq. **B5**) leads to a quantitative description of supersaturation in spiking networks, we generated LIF network parameters fulfilling the supersaturating condition using the SSN-LIF mapping framework in Eq. **1** (Fig. **2A**). We found that the activity in a LIF spiking network and its self-consistency approximation Φ_*sc*_ (Eq. **12**) align robustly with the activity of the SSN model (Fig. **2A**).

**FIG. 2.**
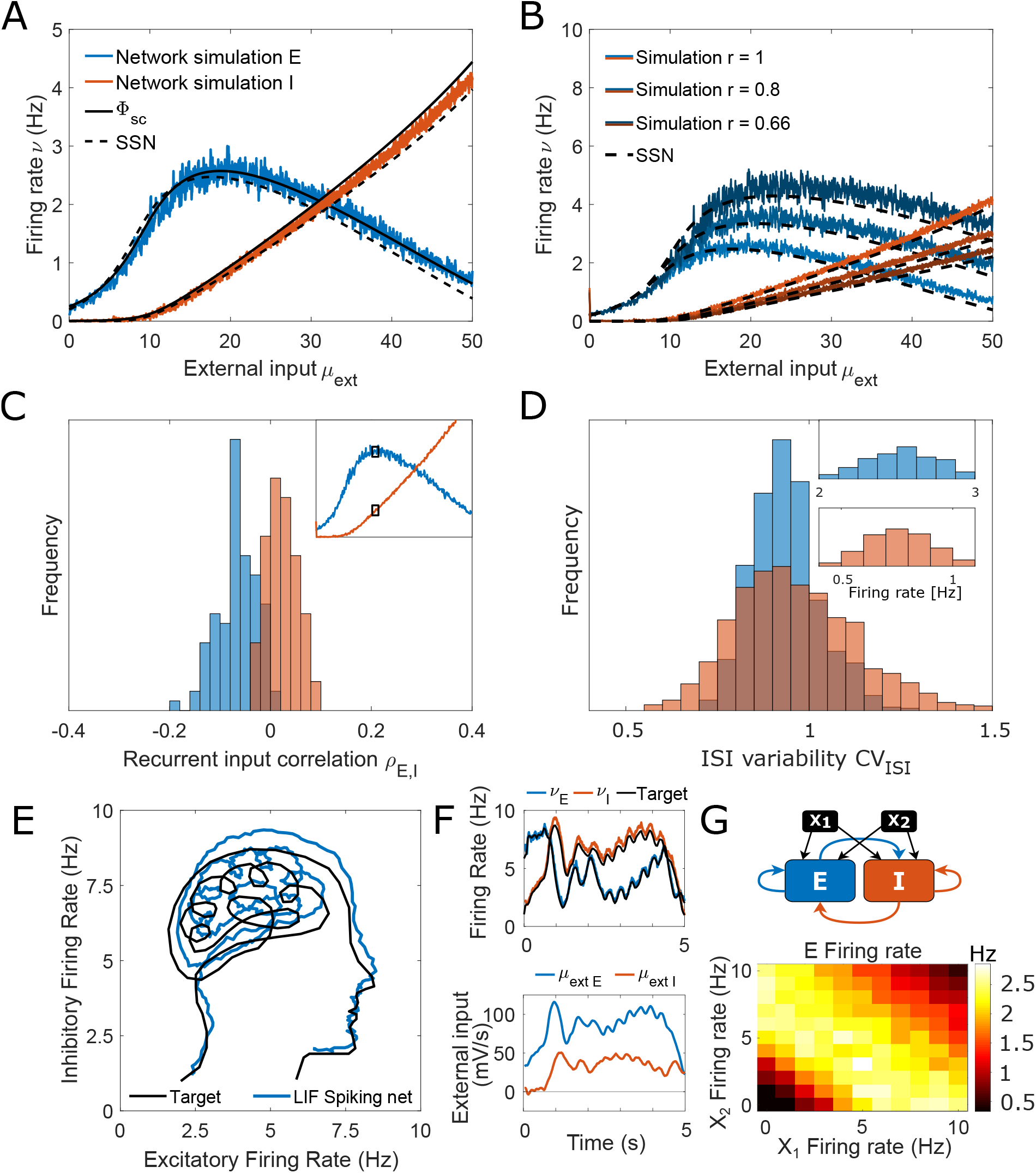
SSN-predicted supersaturation can be observed in spiking network simulations. The existence of the supersaturating response can be predicted by the SSN framework using Eq. **B5**. (A) E (blue) and I (red) firing rates as a function of external input. LIF spiking activity is in line with the self-consistency solution (Φ_*sc*_ in Eq. **12**, solid line) and is also accurately captured by the SSN solution (dashed line). (B) The peak of E activity can be tuned to any desired level by tuning the E/I ratio of external inputs (*r*) along with the connectivity weights *J*_*IE*_ and *J*_*EI*_. (C) Histogram of the E/I input correlation (*ρ*_*E,I*_) onto E (blue) and I (red) neurons computed at the peak of E activity (see inset). The weak correlation suggests the network is operating far from E/I balance. (D) The spiking activity of both E and I neurons is irregular and compatible with a Poisson process (*CV*_*ISI*_ close to 1). The inset shows the distribution of firing rates of individual neurons. (E) Spiking networks can follow a user-defined target dynamical trajectory. The black line shows the target trajectory we aim to replicate with the network. The blue line shows the trajectory of the spiking network in the E-I activity phase space. (F) Same simulation as in panel E, The time course of the E and I firing rates in the LIF network (top) follows the target trajectory and results from designed dynamical inputs (bottom). (G) Supersaturating networks can perform the XOR task. top: Layout of the network used to perform the XOR task, where the E-I network is supersaturating (Eq. **4**). Bottom: The LIF E population activity performs XOR logical operation of the two inputs *X*_1_ and *X*_2_. The feedforward weights are 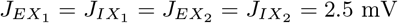. The spiking network parameters can be found in Table **II**.

Recent work [27] compared the responses of LIF and SSN models, pointing out that the peak E activity in supersaturating spiking networks is small, and in particular, it is smaller than the SSN peak. As shown in Fig. **2A**, the peak firing rates obtained with the two methods are in agreement. Furthermore, we show that it is possible to control the height of the E firing rate peak in both networks such that it can be made arbitrarily high (Fig. **2B**, Appendix). Specifically, we show that the peak of E activity can be controlled by modifying the ratio of the external inputs *r*, and the connectivity parameters *J*_*EI*_ and 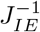 by the same factor. This manipulation derived from the SSN analysis (Appendix) leads to the same effect in the spiking networks (Fig. **2B**).

To determine how close the network operates to E-I balance, we measure the Pearson correlation of the time series of recurrent excitatory and inhibitory inputs to each neuron *ρ*_*E,I*_. The E-I correlation measured at the peak E firing rate is close to zero, demonstrating that the network operates far from E-I balance (Fig. **2C**). Since the coefficient of variation of the interspike intervals at the peak E firing rate is close to 1 (*CV*_*ISI*_ ≈ 1), the firing appears to be irregular, asynchronous, and compatible with a Poisson spiking process (Fig. **2D**). Importantly, the supersaturation regime occupies the biologically plausible activity range of 0-10 Hz in spiking networks [30–37], and the amplitude of the synaptic connection strengths, as well as the size of the network (*N* = 4000), are both in line with biological estimates of functional cortical network size [6, 53, 54] (Appendix). We note that the supersaturation condition is incompatible with the existence of a balanced state solution, as defined in Eq. **B1** as it would lead to negative firing rates.

Knowing how the 2D firing rates emerge from recurrent and feedforward connectivity in the SSN allows us to invert this relation and select external inputs such that they lead to the desired E and I activity trajectory in the spiking network. This is illustrated in Fig. **2E** where we targeted a complex 2D trajectory. We obtained the feed-forward inputs that result in the desired dynamics *ν*_*E*_(*t*) and *ν*_*I*_ (*t*), by inverting Eq. **3**:

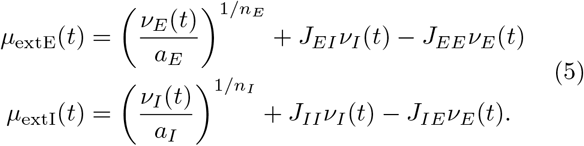

These dynamic feedforward inputs *µ*_extE_(*t*) and *µ*_extI_(*t*) are shown in Fig. **2F, bottom** and the fidelity of the targeting is illustrated in Fig. **2F, top**. Notably, the timescale of the autocorrelation function of neuronal activity (as defined in [55]) is around 300 ms, which is in line with recorded cortical activity [44–46]. These results indicate that complex dynamic trajectories evolving on biologically realistic timescales can be accurately captured by the SSN steady states Eq. **3**.

Let us note that while we used here dynamic feedforward inputs to move along the activity trajectory, it is equally possible to dynamically modify the connectivity to obtain the same 2D trajectory in activity space. In this scenario, synaptic plasticity is recruited to obtain a user-defined output. This can be done by setting the plastic connections *J* as dynamic while the external inputs are constant.

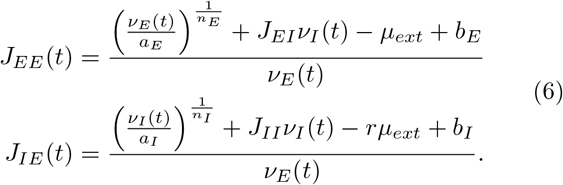

Overall, we show that the mapping between SSN and spiking networks makes it possible to construct inputs or synaptic weights in a spiking neural circuit such that its activity follows a user-defined complex target dynamical trajectory..

In balanced networks, the implementation of logical gates is a complex task due to the linearity of the transfer function [23]. Therefore, we asked whether the nonlinear regimes of spiking networks can be used to perform specific logical operations. Here, we show that it is possible to combine feedforward and recurrent inputs in a way that makes the circuits perform the nonlinear XOR operation, which is one of the key computing components of logical circuits, while being challenging to implement in a neural network [56]. We show in Fig. **2G** how a supersaturating network can execute the XOR operation from two input signals. The E activity is maximal if the input *X*_1_ + *X*_2_ corresponds to the peak input in the SSN supersaturating regime. The E activity is unstimulated if both inputs *X*_1_ and *X*_2_ are low and silenced if they are both high. This shows that the nonlinearity of medium-sized spiking networks can be exploited to carry out fundamental logical operations.

Next, we explore if the experimentally reported connectivity parameters in mouse V1 by the Allen Institute [6] are consistent with supersaturation (Appendix, Table **II**). We use these parameters as a reference point for the biologically plausible range of connection strengths. Interestingly, a circuit with these connectivity parameters does not have a balanced state solution for the equal external input ratio *r* = 1 (*µ*_extE_ = *µ*_extI_) and requires *r <* 0.9 to fulfill the balanced state requirement (Eq. **B1**). The network can be supersaturating for values of *r* larger than 0.9. Remarkably, for *r* = 1, the E activity does not decrease in the low input range but saturates instead (Fig. **3A**), the network only becomes supersaturating for *µ*_ext_ *>* 150 mV/s (Fig. **1B**). For larger values of *r* the activity decreases and becomes silent for inputs close to 100 mV/s (see inset Fig. **3A**, *r* = 1.5).

**FIG. 3.**
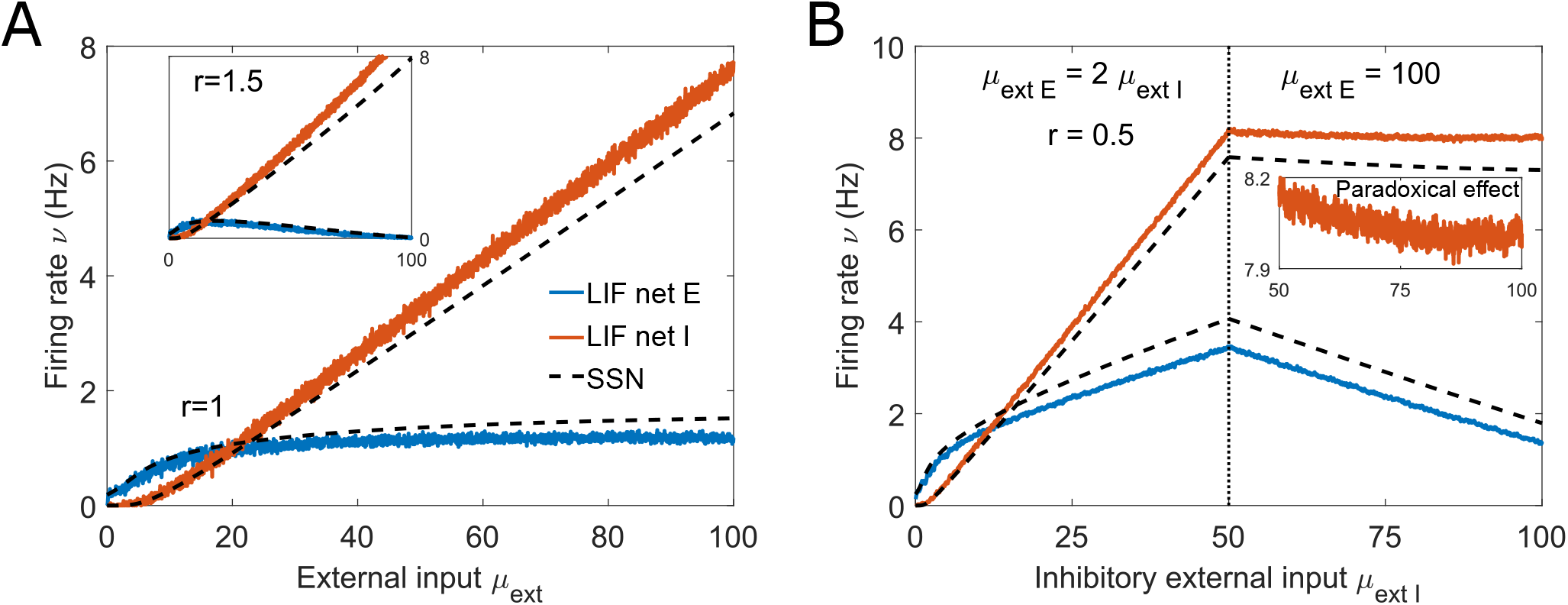
The experimentally reported network parameters can generate supersaturation and be adapted to enter the inhibition-stabilized state. (A) Firing rates of the E and I populations, blue and red lines, respectively, as a function of external input using the parameters reported by the Allen institute [6], see Table **II**. The dashed lines show the SSN solution. In this network, the E activity saturates for inputs larger than 20 mV/s. If the external input to the I population is larger than that to the E population, as shown in the inset with *r* = 1.5, E firing rate declines for growing input. (B) This spiking network exhibits inhibitory stabilization matching the predictions of the SSN (dashed line). The connection strength *JEE* is higher than in panel A (Table **II**). The ISN state is exposed by the paradoxical effect which occurs when the I firing rate decreases for increasing *µ*_extI_. First, the inputs to both populations grow to drive the network in a state where the E subnetwork is unstable: *ν*_*E*_ *>* 1.5 Hz, Eq. **7** (vertical dotted line). Once *µ*_extI_ reaches 50 mV/s, only the input to I increases (from 50 to 100 mV/s), while *µ*_extE_ remains at 100 mV/s. This results in a decrease in the firing rates of both populations, as predicted by the SSN (dashed lines). The inset shows a close up of the I activity to illustrate the paradoxical effect and shows that the paradoxical effect wanes as the E firing rate approaches the ISN threshold (*ν*_*E*_ *≈* 1.5 Hz).

**TABLE I.** Spiking network parameters used in all figures by default. Parameters deviating from the default ones are specified in figures’ panels and captions. For derivation, see section *Extraction of experimentally reported network parameters*.

|  |  |  |
| --- | --- | --- |
| $N_E$ | 3000 | |
| $N_I$ | 1000 | |
| $p_{EE}$ | 6.5 | % |
| $p_{EI}$ | 20 | % |
| $p_{IE}$ | 27.5 | % |
| $p_{II}$ | 10 | % |
| $V_r$ | 0 | mV |
| $\theta$ | 1 | mV |
| $\tau_E$ | 20 | ms |
| $\tau_I$ | 10 | ms |
| $\sigma_{ext}$ | 3 | mV/$\sqrt{s}$ |
| $r$ | 1 | |

**TABLE II.** Network connectivity parameters used in all figures. The synaptic weights *j*_*XY*_ correspond to the strength of a single spike in spiking networks and are given in *µ*V. The connection strength at the population level *J*_*XY*_, used in mean field solutions, are given in mV (Eq. **13**). For Fig. **S2** and Fig. **6**, synaptic weights are N-dependant since the 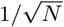 scaling is applied; the given weights correspond to *N* = 4000. See section *Extraction of experimentally reported network parameters*.

| | $j_{EE}$ | $j_{EI}$ | $j_{IE}$ | $j_{II}$ | $J_{EE}$ | $J_{EI}$ | $J_{IE}$ | $J_{II}$ |
| --- | --- | --- | --- | --- | --- | --- | --- | --- |
| Mouse V1<br>Fig. 3A<br>Fig. S1B | 3.59 | 68.1 | 28.7 | 108 | 0.672 | 13.2 | 23.7 | 11.8 |
| Fig. 2<br>Fig. S1A<br>Fig. 6A-B | 10.3 | 60 | 7.27 | 10 | 2 | 12 | 6 | 1 |
| Fig. 4<br>Fig. 5A<br>Fig. S2<br>Fig. 6C-D | 25.6 | 50 | 8.49 | 110 | 5 | 10 | 7 | 11 |
| Fig. 5B<br>Fig. 6E-F | 6.41 | -5 | 1.21 | -12.5 | 1.25 | 1 | 1 | 1.25 |
| Fig. 3B | 25.4 | 68.1 | 28.7 | 108 | 4.75 | 13.2 | 23.7 | 11.8 |

### C. Inhibitory stabilization and its presence for reconstructed synaptic weights

Inhibitory stabilization is a network state in which the recurrent excitation feedback loop is strong and intrinsically unstable but can be stabilized by the recurrent inhibition [25, 57]. The paradoxical effect is a feature of the ISN [25], in which the I activity decreases as the input to the I population is increased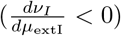. Recent studies using optogenetic stimulation of inhibitory neurons confirmed the paradoxical effect in mouse visual, somatosensory, and motor cortices [58] suggesting that the ISN is a ubiquitous property present across cortical networks. A recent review presented further experimental evidence and techniques used to study the inhibition-stabilized dynamics and discussed the ISN consequences for cortical computation [29]. In the SSN model [47, 48], a network is inhibition-stabilized if it fulfills the condition

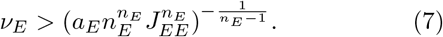

We note that in networks with a threshold linear transfer function, the analogous ISN condition only requires a strong recurrent coupling *J*_*EE*_ *>* 1 and does not impose any constraints on the E firing rate level or the transfer function parameters [25, 29]. However, large enough *J*_*EE*_ does not always guarantee that a recurrent neural network with a nonlinear transfer function is in the ISN regime. Increasing *J*_*EE*_ can also lead to instability, as the excitatory feedback loop can strengthen to a point where it escapes stabilization from recurrent inhibition. In the extreme case, it is even possible to build a network that can never enter the ISN regime regardless of the value of *J*_*EE*_, as E activity never reaches the level where it can be stabilized by inhibition (Appendix).

Next, we investigated whether this condition (Eq. **7**) can predict the existence of the ISN in spiking networks of LIF neurons. Interestingly, we found that the ISN condition cannot be met for the connectivity strengths reported for mouse V1 from the Allen Atlas [6] if the E/I input ratio *r* is equal to 1. This is due to the fact that the required E firing rate (*ν*_*E*_ *>*27 Hz) is higher than the maximal possible stable E firing rate in the network (Fig. **3A**). For very low values of *r* (around *r* = 0.1), an ISN state can only be reached by exposing the network to very high external inputs (around *µ*_ext_ = 1000 mV/s) (Fig. **1B**). We will choose a network which operates outside these cases since the corresponding firing rates are far beyond the 0-10 Hz firing rate range we consider in this study. Therefore, to meet the ISN condition, we modified one of the connectivity strengths. Specifically, we increased the connectivity parameter *J*_*EE*_, which is supported by the study by [10] who report larger *J*_*EE*_ than the Allen Atlas [6]. We chose *J*_*EE*_ such that the network is in the ISN state for E firing rates larger than 1.5 Hz and kept all other connectivity strengths as reported by the Allen Atlas ([6], see Table **II**). Fig. **3B** shows that the resulting network exhibits the paradoxical effect and is therefore in the ISN regime.

### D. Bistability and persistent activity

One of the most prominent experimentally recorded neural activity features *in vivo* is the network ability to switch between higher and lower firing levels. One example is spontaneously alternating intervals of tonic firing and silence observed across different cortical areas [34]. Another example is the sustained firing rate in the prefrontal cortex after stimulus withdrawal during decision-making tasks which is hypothesized to represent short-term memory [14, 59]. The coexistence of multiple network states can be explained theoretically by bistability, where the system has two stable states for the same level of input. If multiple stable states co-exist in a network model, a sufficiently large perturbation can drive network activity away from its current state towards another attractor. In the situation where a bistable network can sustain its high activity level in the absence of feedforward input, it has persistent activity. Here, we asked whether the SSN model can predict the connectivity regime supporting bistability in spiking circuits.

Bistability and persistent activity can be obtained in the SSN model [28] without the need for synaptic plasticity [21] or complex synaptic weight distributions [22]. Unlike supersaturation and inhibition stabilization, bistability cannot be delimited by a simple tractable condition on network parameters (Appendix). However, we can use the conditions presented in [28], as a starting point to guide our search for bistability in biologically realistic spiking networks, even though they are derived under restrictive assumptions on the *a, b*, and *n* parameters. We show an example of a biologically realistic bistable network in Fig. **4**.

**FIG. 4.**
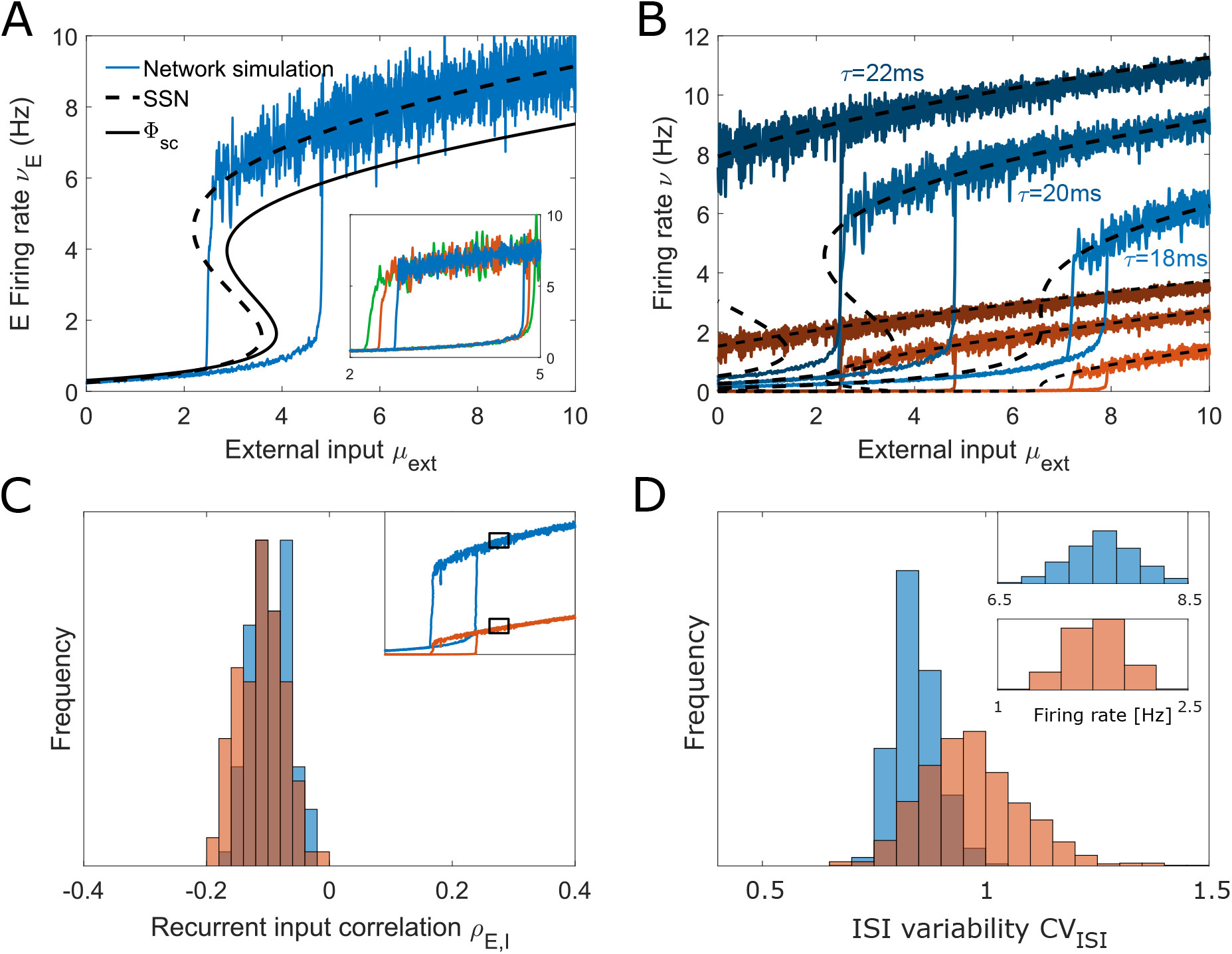
SSN-predicted bistability and persistent activity can be observed in spiking network simulations. (A) E and I firing rates as a function of external input in a bistable network (coexistence of high and low activity states for a given external input). Simulated LIF spiking activity is in line with the self-consistent solution (Φ_*sc*_) and is also accurately predicted by the SSN. The Φ_*sc*_-predicted firing rates diverge slightly from the spiking network simulation because of the use of exponential synapses, which lead to correlated recurrent noise (Methods, Appendix). The inset illustrates that the width of the bistability window can vary between simulations of the same network due to the spontaneous transitions between the two states. (B) The width of the bistability window depends on the excitatory membrane time constant: higher values of *τ*_*E*_ lead to broader bistability windows, which are shifted leftward. If *τ*_*E*_ is sufficiently large for the bistability window to exist for zero feedforward input, the network can sustain persistent activity. (C) E/I input correlations to E (blue) and I (red) neurons indicate a persistent but weak E/I correlation, suggesting that the network is only loosely E/I balanced. (D) The coefficient of variation of the interspike intervals (*CV*_*ISI*_) is near 1, which is compatible with a Poisson process and demonstrates that activity is asynchronous and irregular. Both (C) and (D) are measured in the upstate, as shown in the inset to (C). All parameters are given in Table **II**.

Both the self-consistency solution and LIF network simulation confirm the SSN-predicted bistability Fig. **4A**: the network can sustain either low activity or high activity for external inputs in the 2-4 mV/s range. Although the SSN and Φ_*sc*_ are deterministic, the spiking network simulation is not. Due to the stochastic nature of the neuronal activity, fluctuations in firing can cause spontaneous transitions between steady states (shown in Fig. **4A**, inset). We note that the spontaneous transitions between the up and down states have not been reported in the bistable balanced networks with short-term plasticity [21] because spontaneous fluctuations in average firing and the probability for spontaneous transitions decrease with network size.

We find that a higher excitatory membrane time constant broadens the window of bistability (Fig. **4B**), making bistability more robust to spontaneous fluctuations and easier to locate in phase space. As *τ*_*E*_ increases, the bistability window shifts towards lower feedforward input. When the bistability window intersects the vertical *µ*_ext_ = 0 axis, the network has a persistent activity state in the absence of feedforward input (*τ*_*E*_=22 ms in Fig. **4B**). Here again, the E-I balance is weak, as shown by the correlation in recurrent inputs (Fig. **4C**), and the spiking activity is Poisson-like, as shown by the coefficient of variation of the ISIs (Fig. **4C**).

Finally, the nonlinear transformation performed by spiking networks can be functionally relevant for information processing. Logical operations such as the AND operation can be implemented without the need to recruit synaptic plasticity, thanks to the sharp transition between the two stable states. If the transition from the low to the high activity level requires a strong input, so that two signals *X*_1_ and *X*_2_ need to be present to elicit the transition, the network can execute the AND operation. Moreover, the bistability of a neural network can also offer the possibility to store information. Once the network has been switched into a different activity state by a strong perturbation, it remains in the same state even after the perturbation withdrawal.

### E. Computational regimes and their position in

In previous sections, we demonstrated that the SSN framework can be used to locate specific computational regimes such as supersaturation and the paradoxical effect in parameter space. Here, we focused on the activity regimes associated with the 2D space of the feedforward input and the I/E external input ratio (*µ*_ext_, *r*). For two examples of connectivity matrices *J*, we scanned the 2D input space for supersaturation (Eq. B5), ISN (Eq. **7**) and bistability using the characteristic function ℱ as defined in [28] (Eq. **B10**). We also show the input regimes for which the network permits a balanced limit solution (Eq. **B3**). Importantly, the sign of the determinant of the weight matrix (det *J* = *J*_*EI*_ *J*_*IE*_ − *J*_*EE*_ *J*_*II*_) determines the number of SSN steady states (stable and unstable): for det *J >* 0, the SSN has an odd number of steady states whereas for det *J <* 0 the number of steady states is even [15, 28]. Thus, networks with positive det *J* (Fig. **5A**) always have at least one steady state. In the network shown in Fig. **4**, bistability occurs when the system transitions from having one steady state to having three (two stable and one unstable). On the other hand, networks which have a negative det *J* (Fig. **5B**) can lack steady states at all, and any possible stable steady state coexists with an unstable steady state [28]. Finally, the sign of det *J* also determines whether the system can have a stable balanced state (Eq. **B3**) or lacks it. In networks where the sign of det *J* is negative, a balanced state solution can exist with positive firing rates if 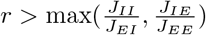 but it is unstable [24].

**FIG. 5.**
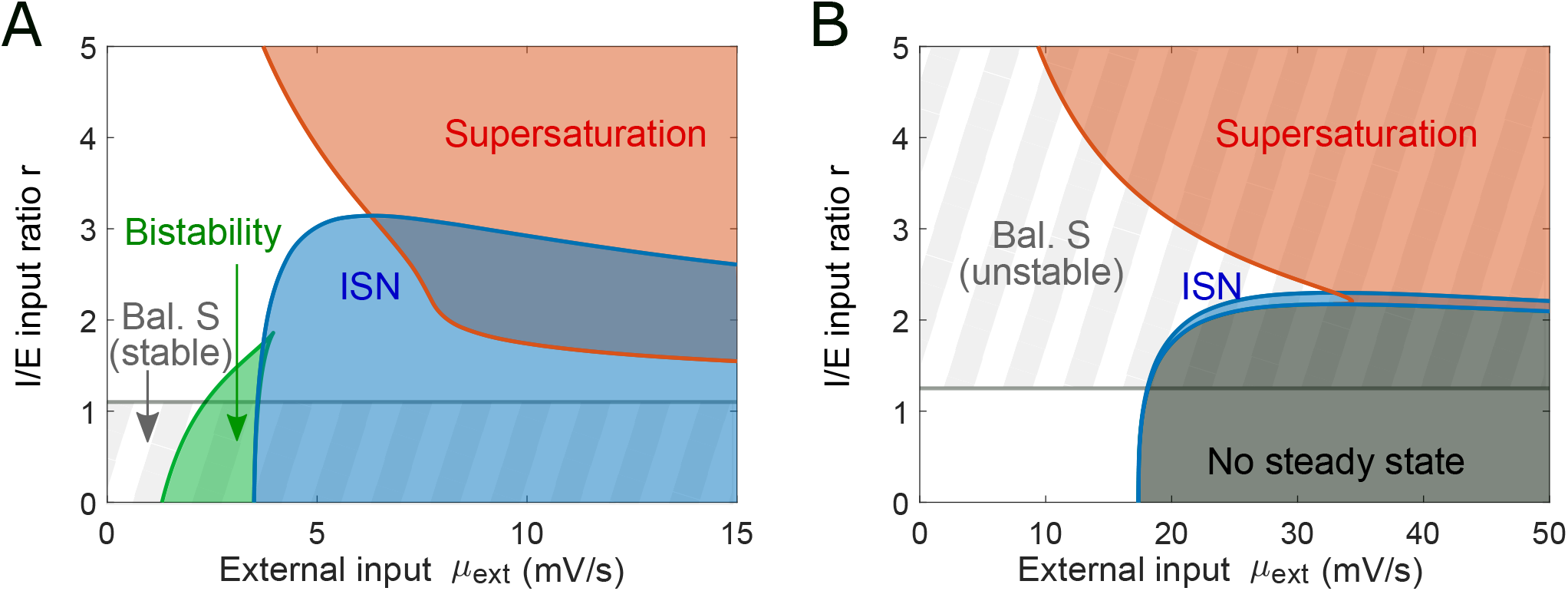
Mapping the computational states in the SSN model for two representative connectivity regimes. (A) Varying the ratio of the external input weights and the amplitude of external drive in a network with a positive det *J* allows to traverse different computational regimes (*J* as in Fig. **4**, see parameters in Table **II**). Gray stripes denote the input space subset with a stable balanced state limit (*N→ ∞*) which does not exist above the horizontal gray line. The blue area represents the inhibition stabilized regime (ISN). The red area denotes the phase space occupied by supersaturating spiking activity. The green area corresponds to a bistable region (as shown in Fig. **4A** with *r*=1). Within the green region, the up-state is in the ISN whereas the down-state is not. We note that the inhibitory stabilization and supersaturation can co-exist. (B) The same analysis is performed on a network with negative det *J* (Table **II**). In this case the balanced state limit is unstable. The ISN region is narrower and there is a broad range of inputs for which the network does not have a steady state solution.

Fig. **5** shows the map of feedforward inputs and the corresponding computational regime for two examples of the connectivity matrix *J*. Panel A corresponds to the connectivity parameters from the bistable network shown in Fig. **4** with det *J >* 0. The region where bistability is expected corresponds to the results in Fig. **4A** with *r* = 1. The balanced state exists and is stable for low values of *r*. Panel B corresponds to a network with det *J <* 0. In this case, the balanced state only exists for high values of *r*, but it is unstable. Furthermore, we find that for large input and low *r*, the network does not have a steady state. In this region, inhibition cannot stabilize the network, and the activity blows up. The same analysis is also performed for the supersaturating network in Fig. **2** and the mouse V1 network in Fig. **3A** (Fig. **S1**). Overall, using the SSN model, we can precisely locate the regions corresponding to distinct behaviors of spiking networks in their parameter space. Notably, we observe that the sign of the determinant of the connectivity matrix *J* plays a crucial role in the type of activity regimes available to the network (Appendix).

### F. Effect of network size on network response nonlinearity

While medium-sized networks can generate diverse nonlinear responses to external input, the balanced state framework implies that network response becomes linear as network size approaches infinity. How do networks transition from nonlinear to linear regimes for increasing network size *N*? To tackle this question, we re-scaled the recurrent connections *J*_*XY*_ by the factor 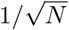 as a function of network size *N*, and increased *N* from *N* = 4 ×10^3^ to 5 ×10^5^. This parameter re-scaling follows the convention of the balanced state theory [2, 16] and allows us to address whether these nonlinear spiking networks converge to the expected balanced state, and if so, when and how.

A dynamically stable balanced state limit can only exist if det *J* is positive and the fraction of external input weights *r* satisfies 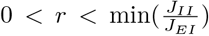, see Eq. **B3**. In our network convergence study, we focus on three spiking networks: one supersaturating network with det *J >* 0 (shown in Fig. **2**), one bistable network with det *J >* 0 (shown in Fig. **4**), and a supersaturating network with det *J <* 0 (presented in Fig. **5B**, where we set *r* = 3). Among our three example networks, we have one example for which a balanced state limit does not exist (supersaturation), one network with a stable balanced state solution (bistable network) and one network with a balanced state that exists but is dynamically unstable (det *J <* 0). In all three cases, the self-consistency solution Φ_*sc*_ remains an accurate description of the spiking network mean activity across different network sizes *N* (Fig. **6A, C, E**), and its predictions align qualitatively and quantitatively with the SSN model (Fig. **6B, D, F**, inset).

**FIG. 6.**
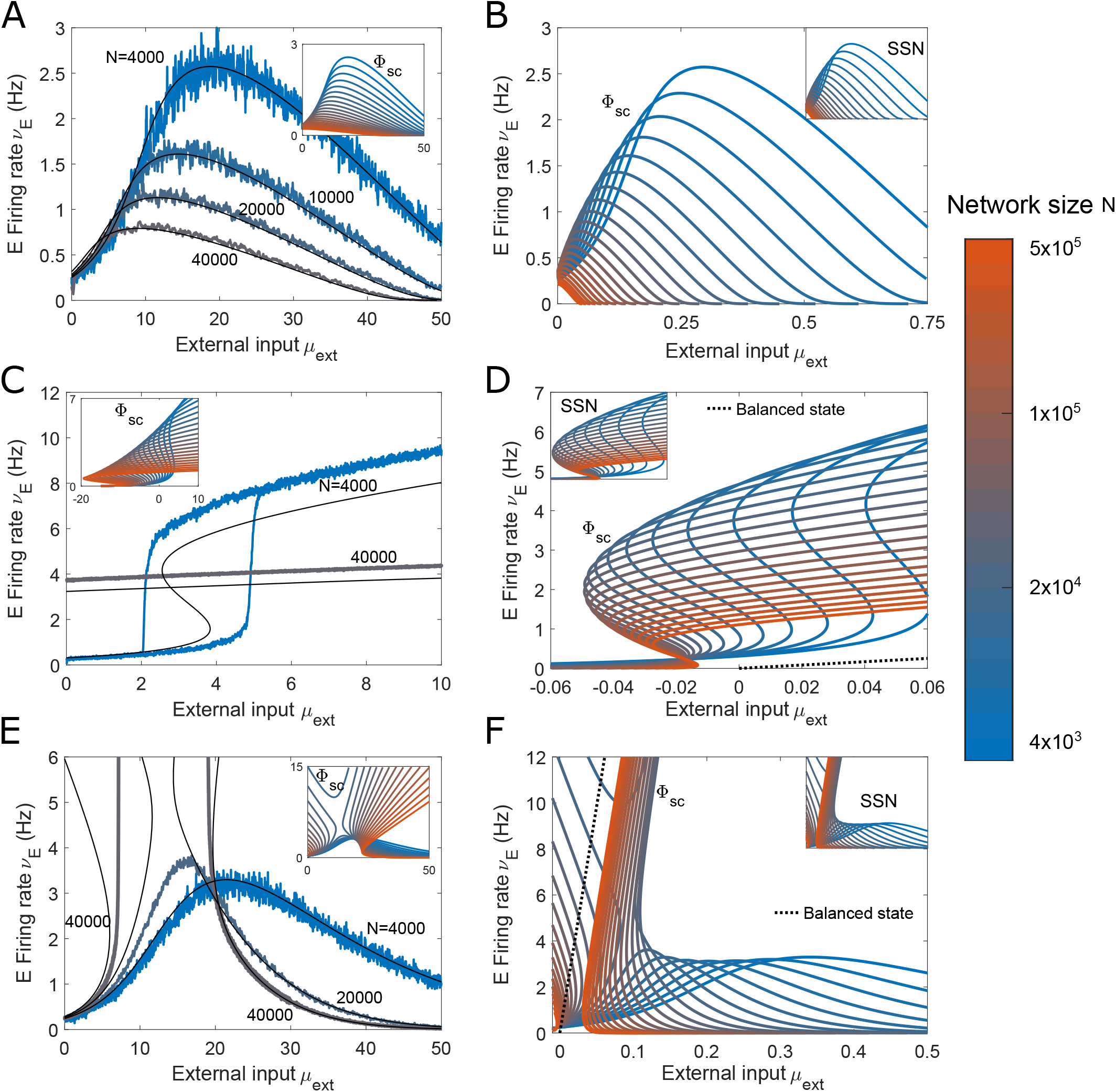
Increasing network size does not guarantee convergence to a balanced state. (A-B) supersaturating network with det *J >* 0, (C-D) bistable network with det *J >* 0, (E-F) supersaturating network with det *J <* 0, parameter regimes which we identified using the SSN framework and studied for *N* = 4000 in previous figures. Here, we gradually increase the size of these networks *N* and follow the balanced network convention to rescale the weights *J*_*ij*_ by 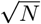 as the network grows. depicts the excitatory firing in the supersaturating network from Fig. **2A** across different network sizes. The colored lines represent spiking networks (from 4 *×* 10^3^ to 4 *×* 10^4^), the black lines represent the corresponding mean-field solution Φ_*sc*_ (Eq. **12**). The inset shows the excitatory rate for network sizes from *N* = 4 *×* 10^3^ (blue) to *N* = 5 *×* 10^5^(red), in steps of *×*10^0.1^ obtained using Φ_*sc*_. (B) depicts the same network as in A, but the external input is rescaled as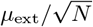. This network does not have a balanced state solution (see first condition of Eq. **B3**). As *N* grows, the excitatory activity peak becomes smaller and in the limit of very large networks, the excitatory population remains silenced (*ν*_*E*_ = 0). The inset shows that the SSN and Φ_*sc*_ predict the same behavior as N increases. (C) shows the excitatory activity for the bistable network from Fig. **4**, for *N* = 4 *×*10^3^ and *N* = 4 *×*10^4^. The spiking activity of the spiking LIF network (colored lines) is captured by Φ_*sc*_ (black lines). As *N* increases we observe a broadening of the bistability window and a decreasing firing rate, see inset. (D) depicts the same network as in panel C but now with rescaled external inputs. The balanced state predicts a linear solution for *N*→ ∞ limit (dashed line). The convergence to the balanced limit is very slow, and even at *N* = 5 *×*10^5^ neurons, the network rates still do not converge to the balanced state. (E) Activity of a spiking network with a negative det *J* from Fig. **5B**, with r=3. As *N* grows, the excitatory activity dissociates into two distinct branches separated by an unstable region where the firing rates diverge to ∞. The inset illustrates that this separation occurs when the stable and unstable steady states collide. (F) depicts the same network as in E, now with rescaled external inputs. The balanced solution of this network (dashed line) is unstable (Eq. **B3**). The unstable solution of Φ_*sc*_ approaches the balanced state, while the stable solution tends to 0 as *N* increases.

We find that the network response can remain nonlinear even for very large network sizes consisting of up to half a million neurons (see inset). By re-scaling the feed-forward input with the factor 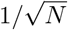 [2, 16], the network response should converge toward a single linear solution - the balanced limit (Fig. **6B, D, E**). In the case of the supersaturating network, the balanced limit does not exist, as it would lead to negative E firing rates. Therefore, in the limit of large network size the E firing rate tends to zero. In the case of the bistable network, a balanced state limit does exist but the network response is still far from converging to it, even for *N* = 5 × 10^5^. Finally, for the network with det *J <* 0 and *r* = 3, the network exhibits supersaturation for *N* = 4000 (see Fig. **5B**). However, as *N* increases, the network enters a region for which there is no steady state, and where the firing rates blow up. This behaviour is observed in spiking network simulations, the mean field Φ_*sc*_ and the SSN solution. Interestingly, the LIF simulation appears to be more stable than the mean-field predictions, and for *N* = 20000 the firing rates grow substantially but do not explode as it is the case for the mean-field solution. The inset shows how this instability is caused by the collision of the two steady state branches, leaving a gap where the firing rates are unbound. For this network, the mean-field solution converges to the balanced limit as *N* increases (Fig. **6F**). However, the balanced state limit is unstable here, and it only matches with the unstable mean-field solution (high firing rates part of the branch) whereas the stable low activity solution tends to zero.

Overall, our example networks illustrate that for many classes of spiking networks with biologically plausible sizes and connectivity configurations, the activity will escape the predictions of the balanced state. Depending on the parameters, the balanced state may not exist or be unstable. Yet, even when a stable balanced state exists, it is not guaranteed that it provides a realistic description of network activity, even for unrealistically large sizes *N*. Next, we investigated whether the correlations between recurrent I and E inputs that accompany the balanced state can also be observed in our networks that are non-linear and do not conform to the balanced predictions. We have shown that the correlation between E and I recurrent inputs *ρ*_*E,I*_ are weak in the bistable network of 4000 neurons (Fig. **4C**), which is an indicator that the network is operating far from balance. However, we found that *ρ*_*E,I*_ can be network size dependent: increasing the size *N* from 4000 to 40000 strongly increased the correlation even though the firing rate activity in both cases does not conform to the balanced state solution (Fig. **S2**). In summary, the observation of strong E-I correlation does not guarantee that the balanced state framework is applicable to predict the firing activity.

## III. DISCUSSION

Understanding the activity regimes of biologically sized spiking networks is critical to make sense of experimentally recorded data. The state-of-the-art experimental techniques now enable simultaneous recordings of thousands of neurons [60, 61]. Therefore, it is important to relate experimentally recorded network activity to the computational capabilities of similarly sized networks *in silico*. On the theoretical side, many commonly used models operate in the limit of infinitely large networks (balanced state) [2, 21–24]. Some features predicted by the balanced state, such as high correlations between excitatory and inhibitory currents within neurons, have been observed experimentally [62, 63]. However, other features, such as strong feedforward inputs, have not been experimentally reported, and some experiments argue against it [22, 64–67]. Unlike the balanced state framework, which assumes a tight E/I input balance, alternative models such as the SSN only assume loose E/I balance to achieve inhibitory stabilization [20, 29, 48, 57, 58]. Yet, clarifying under which conditions the predictions of these two models are conflicting or aligning and how they compare to spiking network simulations has so far remained an open question.

Here, we mapped the computational regimes of spiking networks containing a few thousand neurons across the input space. We chose this size because neuroanatomical and connectomics studies of different cortical regions such as the somatosensory cortex [68–72] indicated that a functional unit such as a minicolumn could contain a few thousand neurons. We also set the range of firing rate activity such that it meets the experimentally reported range of a few Hz [30–37].

We have shown that the nonlinear behavior of medium-sized spiking networks can be quantitatively and qualitatively understood using the SSN model which is based on the power-law activation function of individual neurons. Importantly, the mapping we propose can address the broad activity and connectivity regimes, including those located outside the validity domain of the balanced network formalism. We delineated connectivity regimes where bistability, supersaturation, inhibitory stabilization, or even the absence of steady states can occur in spiking networks. We found that networks can be inhibition stabilized in conditions where a balanced limit does not exist, even though both require strong inhibitory feedback. The ISN can overlap with supersaturation or can be achieved in networks with det *J <* 0 (Fig. **5**), which is outside of the domain of the balanced state framework. Furthermore, we found that network parameters obtained from experimental recordings are incompatible with a balanced solution unless the E population receives a stronger feedforward input than the I population.

We used our theory to target a 2D firing rate trajectory of interest within the spiking neural circuit and could implement an XOR gate by exploiting the intrinsic non-linearity of the neuronal transfer function (Fig. **2E,G**). This suggests that designing a biologically realistic spiking network to perform a task can be facilitated by understanding the input-output relationship in networks.

Studying the network responses across different network sizes, we found that convergence to the balanced state was not always guaranteed, even for unrealistically large network sizes. Activity deviations from the predicted balanced solutions were significant and could be observed even for large E/I correlations of the recurrent inputs. This indicates that strong E/I correlations can be present in networks that do not meet the balanced network predictions on the activity level (e.g., exhibit bistability, Fig. **S2**). At the same time, networks can be asynchronous irregular in the absence of strong E/I correlations (e.g., Fig. **2**). Importantly, we showed that the SSN model could accurately describe spiking networks that do not have a stable balanced limit.

Overall, the balanced state framework has several limitations such as a limited domain of validity, a linear response function, the need for extensive network sizes, and strong feedforward inputs. We show that we can avoid these limitations by using a different rate model, the SSN which assumes a power law transfer function. As we show here, these limitations can be addressed in medium-sized spiking networks by mapping them to the SSN model, which supports nonlinear responses for a broad range of connectivity configurations without the requirement of strong feedforward inputs. It should be noted, however, that other works are addressing these limitations within the balanced state framework, by expanding it. For example, balanced networks with shortterm synaptic plasticity have been proposed to permit the emergence of nonlinear activity, such as bistability [21]. Likewise, the experimentally reported small feedforward input which drives spiking activity *in vivo* [64–67] was inconsistent with the original balanced state predictions but was accommodated via the inclusion of broad synaptic weight distributions [22]. Similarly, semi-balanced networks were proposed [23], where neurons which receive net inhibition remain silent. This generated a piecewise-linear manifold which can operate as a nonlinear decision boundary and allowed for a broader domain of validity than the classical balanced framework. Let us also mention that [27] considered a neuronal transfer function which is nonlinear at onset (noise driven), saturating at high rates (due to a refractory period) and linear in between. This study showed quantitative differences in a set of nonlinear responses of SSN and spiking network models which we resolved using a precise mapping between SSN solutions and spiking network activity.

Our modeling approach considers two homogeneous neural populations, which operate at equilibrium. This approach led to a model with tractable equations and only a few parameters. The complementary modeling strategy, which aims to provide a highly detailed description of a neural system [73] achieves precise biological realism at the expense of mathematical tractability. Future studies could expand the results presented here by including additional features into the network and studying their impact. Such features could include synaptic plasticity which would provide an additional source of network nonlinearity and lead to an even richer repertoire of operational regimes. Similarly, future work could analyze the dynamical properties of spiking networks, as they have been shown to affect the stability of fixed points or to be linked with activity regimes such as oscillations observed in the SSN model [28] (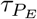 and 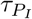 in Eq. **2**). Moreover, the distribution of synaptic weights has been shown to play a significant role in the overall activity regime of spiking networks [22, 74], suggesting that heterogeneity in the properties of neuronal populations could be an important feature of spiking networks which future works could include in the SSN prediction. Finally, our study of medium-sized spiking networks (∼ 10^3^ neurons) lays the groundwork for the analysis of much larger networks, such as a whole functional area (∼ 10^5^ neurons [75]). To that end, the large network would be broken down into a “network of networks” where each system unit represents a single E-I network described by the quantitative mapping developed here.

## IV. METHODS

### A. Power-law approximation of the input-output transformation in a single neuron

We represent the spiking activity of a neuron using the integrate-and-fire model

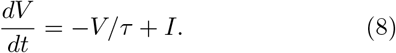

Where *V* is the membrane potential, *τ* is the membrane time constant, and *I* is the input to the neuron. Upon reaching the firing threshold Θ, *V* (*t*) is reset to *V*_*R*_. If we assume the input *I* to be white noise with a mean *µ* and variance *σ*^2^, the firing rate of the neuron in Eq. **8** can be described by the Ricciardi function Φ [39, 76]

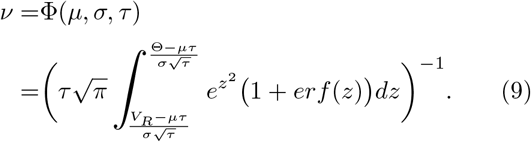

For low inputs, Φ is a supralinear function of the mean input *µ* and can be accurately approximated by a power law with an exponent *n >* 1 (see Fig. **1**). For high inputs, however, Φ becomes linear. In this work, we restrict ourselves to the low firing rate regime (*ν* ≤ 10 Hz) often reported for cortical activity measured *in vivo* [30–37]. In this low activity regime with a constant variance *σ* and time constant *τ*, the firing rate can be accurately approximated by a power law Eq. **1**, see Fig. **1B**. The power law parameters *a, b* and *n* are obtained by fitting Eq. **9** (Fig. **1C**).

### B. LIF spiking network

We consider a spiking network of one excitatory (E) and one inhibitory (I) population with 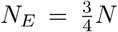 and 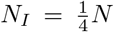 neurons, respectively. We assume that both E and I populations are homogeneous, i.e. neurons within each population have the same parameters (membrane time constant *τ*_*X*_, threshold potential Θ_*X*_, reset value *V*_*RX*_), receive external input with the same mean *µ*_extX_ and variance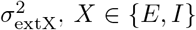. The E and I populations have different membrane time constants (see black crosses in Fig. **1B**), and the feedforward input they receive differs by a factor of *r* (*µ* _extI_ = *rµ*_extE_, see Fig. **1A**). Additionally to the feedforward input, the neurons receive recurrent input from other E and I neurons in the network. The connections are randomly generated based on a homogeneous probability of connection, such that each neuron in population *X* receives inputs from exactly *N*_*Y*_ *p*_*XY*_ randomly chosen neurons in population *Y*, where *p*_*XY*_ is the connection probability from population *Y* to population *X*. We use two types of synapses, the delta synapse and the exponential synapse.

For delta-synapses, the function

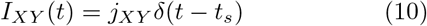

represents the input from a neuron of the population *Y* to a neuron in *X*. Where *j*_*XY*_ is the strength of the synapse, *t*_*s*_ is the spike time of the presynaptic neuron, and *δ* is the Dirac delta function.

In some network configuration, delta synapses promote synchronization of the whole neuronal population. This synchronicity can lead to population spikes [77, 78] which violates the assumption of asynchrony and irregularity in the mean field approach. In order to avoid this synchronization in these cases, we use exponential synapses instead of delta synapses. In exponential synapses, the synaptic potential from a neuron in population *Y* to a neuron in population *X* decays exponentially in time

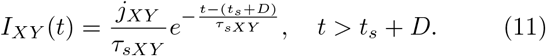

Where *j*_*XY*_ is the strength of the synapse, *t*_*s*_ is the spike time of the presynaptic neuron, *τ*_*sXY*_ is the synaptic decay time constant and *D* is the synaptic delay. This type of synapse prevents synchronization as the effect of each spike is more distributed in time and each synaptic connection has a different delay *D*.

We use the exponential synapse in the spiking network simulation in Fig. **4**, Fig. **6C**, Fig. **6E** and Fig. **S2**, and delta synapses in all other cases.

### C. Self-consistent network solutions

In this work we derive predictions for the activity regimes of spiking networks using the closed-form solutions offered by the SSN framework (Eq. **3**). In some instances, it is useful to compare the SSN predictions to the previously proposed self-consistent network solutions to understand the dynamic origin of the SSN predictions. We provide here the system of self-consistent mean field network equations that arise from the Φ transfer function [78] and that need to be solved numerically to obtain E and I firing rates *ν*_*E*_ and *ν*_*I*_

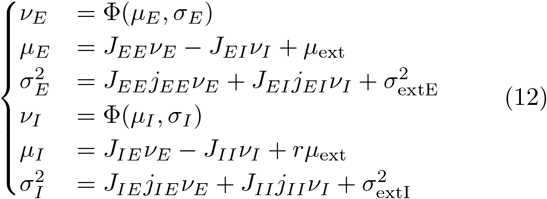

Where *J*_*XY*_ is the population-wide connectivity defined by

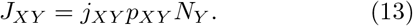

In these equations, the steady state spiking activity of neurons is assumed to follow a Poisson process with a constant rate *ν*_*X*_, *X* ∈ {*E, I*}. The input to each neuron is modeled as a Gaussian process with a mean *µ*_*X*_, and a variance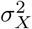. We refer to this approach as “Self-consistency solution” or Φ_*sc*_ in our figures.

### D. Mapping LIF network - SSN

To meet the SSN activity regime with a simulation of spiking LIF neurons, we map the LIF network parameters to SSN parameters. The connectivity parameters *J*_*XY*_ in the SSN correspond to the population-wide connectivity defined for the self-consistency solution Φ_*sc*_ according to Eq. **13**. The transfer function parameters *a, b* and *n* for each of the populations are obtained by fitting the F-I curve of the neuron obtained with the Φ function Eq. **9**, which depends on the LIF membrane time constant *τ*, reset potential *V*_*R*_, firing threshold Θ, and the input noise *σ* (Fig. **1**). The noise *σ* is set to be the external noise *σ*_*ext*_.

In LIF spiking networks, *σ*_*ext*_ models the fluctuations in the membrane potential, which can be caused by fluctuations in the external network input as well as originate from intrinsical properties of the neuron [79]. We note that unlike in the Ricciardi mean-field solution Φ_*sc*_ (Eq. **12**), the SSN framework (Eq. **3**) does not explicitly model the input noise *σ* to neurons embedded in a network. Instead, the effect of the noise is implicitly included in the power law approximation of the F-I curve. As a result, the noise in the SSN model is independent of the network activity leading to the assumption that the noise associated with recurrent input is negligible compared to the external noise 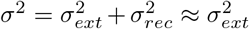. This approximation holds if the firing rates *ν* and the connection strength *j*_*XY*_ are kept low (Eq. **12**), which is supported by experimental evidence [6, 30–37] (Appendix).

## V. ACKNOWLEDGEMENT

This work was supported by the Max Planck Society, University of Bonn Medical Center, University of Mainz Medical Center, German Research Foundation (CRC 1080 to T.T.) and the Loewe Center for Multiscale Modelling in Life Sciences. We thank Laura Bernáez Timón for comments on earlier versions of the manuscript. We thank Laura Busse, Julijana Gjorgjieva, Jochen Triesch, Simon Renner and all members of our group for scientific discussions. We thank Alexandra Vormberg and Andreas Nold for their contribution at the early stage of the project.

## Appendix A: Extraction of experimentally reported network parameters

We use the Cell Types and the Synaptic Physiology databases from the Allen institute [6], to derive biologically plausible network parameters based on the data collected for layer 2/3 of the mouse visual cortex. Nevertheless, our analyses are not restricted to this brain region, as our framework is applicable to any cortical network. It should be noted that the reported values of network parameters vary largely between sources [8–12]. Furthermore, detailed network simulations using state-of-the-art experimentally measured parameters require optimization of all their recurrent connection weights in order to generate realistic spiking activity [73, 80]. Therefore, we use the parameters for mouse V1 extracted from the Allen institute database [6] as a starting point from which we can explore the range of biologically plausible network connectivity.

We use the Cell Type database [6], to obtain the membrane time constant (*τ*) as well as the membrane resting and threshold potentials (*V*_*R*_ and Θ), for E and I neurons. The data has been obtained through whole cell patch clamp recording. We use the Synaptic Physiology database [6] to derive the synaptic strength and the probability of connection (*j*_*XY*_ and *p*_*XY*_) between E and I neurons. The data has been obtained with octopatching, the simultaneous patch-clamp recording of up to eight neurons. The neurons whose dendrite type is classified as *spiny* constitute the E population. The I population consists of all neurons classified as Vip, PV and Sst.

The reported neuronal properties (*τ, V*_*R*_ and Θ) were obtained by averaging the recorded values from 66 E neurons and 94 I neurons. The probability of connection (*p*_*XY*_) is the fraction of connected pairs with respect to all probed pairs, where 80 EE, 150 EI, 160 IE and 607 II cell pairs were probed, Table **I**. Finally, we derive the connection strength (*j*_*XY*_) using the peak amplitude (*A*_*P SP*_), the rise time (*t*_*R*_) and the decay time constant (*τ*_*D*_) of the postsynaptic potential. These parameters are measured over 66 EE, 46 EI, 3 IE and 29 II connected cell pairs. The mean-field analysis of the network at equilibrium does not depend on the dynamical properties of the synaptic transmission (i.e. the shape of the PSP profile). Instead, the mean-field strength of the synaptic connection is the overall depolarization caused by a single presynaptic spike and it is characterized by a single value *j*_*XY*_, regardless of the dynamics. It is given by the time integral of the synaptic current elicited by one presynaptic spike. We assume that the profile of the postsynaptic membrane potential following a presynaptic spike is a linear increase followed by an exponential decay and that the recorded postsynaptic neuron was at resting potential before receiving its input.

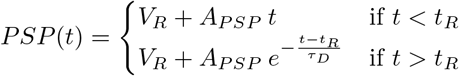

Using the LIF equation (Eq. **8**) for a single presynaptic spike (*Idt* = *j*_*XY*_) yields

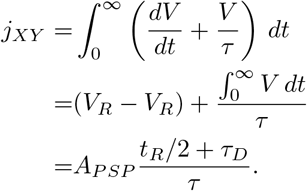

The synaptic strength is then normalized by the experimentally recorded value of Θ − *V*_*R*_ [6], so that *V*_*R*_ and Θ can be set to 0 and 1, respectively.

By default, we assume that both populations receive the same external input (*r* = 1). Finally, we assume the input noise *σ*_*ext*_ to be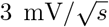, which leads to a power law exponent in the F-I curve close to 3, similar to the values reported in [35, 40] (Table **III** - E and I populations).

**TABLE III.** SSN parameters used in all figures. The parameters *a, b*, and *n* are based on the power-law fit of the LIF F-I curve (Eq. **1**), with the input *µ* in mV/s and the firing rate *ν* in Hz.

| SSN parameters | a | b | n |
| --- | --- | --- | --- |
| Excitatory population | $1.08 \times 10^{-4}$ | -11.1 | 3.08 |
| Inhibitory population | $2.21 \times 10^{-6}$ | 4.8 | 3.82 |
| Fig. 1 ($\sigma = 1 \text{ mV}/\sqrt{\text{s}}, \tau = 20 \text{ ms}$) | $3.91 \times 10^{-2}$ | 31.6 | 2.15 |
| Fig. 1 ($\sigma = 3 \text{ mV}/\sqrt{\text{s}}, \tau = 15 \text{ ms}$) | $3.23 \times 10^{-5}$ | -6.4 | 3.31 |
| Fig. 1 ($\sigma = 5 \text{ mV}/\sqrt{\text{s}}, \tau = 10 \text{ ms}$) | $4.90 \times 10^{-8}$ | -63.0 | 4.20 |
| Fig. 4B ($\sigma = 3 \text{ mV}/\sqrt{\text{s}}, \tau = 18 \text{ ms}$) | $7.89 \times 10^{-5}$ | -9.5 | 3.14 |
| Fig. 4B ($\sigma = 3 \text{ mV}/\sqrt{\text{s}}, \tau = 22 \text{ ms}$) | $1.60 \times 10^{-4}$ | -12.1 | 3.01 |

The neuronal parameters used in LIF spiking network simulations are presented in Table **I**. Fig. **3A** is generated with the mouse V1 parameters we extracted from the Allen institute database [6]. We explore the range of biologically plausible network connectivity parameters by modifying the individual connection strengths *J*_*XY*_ while keeping them within the range delimited by the lowest and largest experimentally reported connectivity values in mouse V1 (0.5≤ *J*_*XY*_ ≤25). The connectivity parameters used in all network simulations are presented in Table **II**.

### Deriving biologically plausible neural network size for circuit simulations

In our framework, we assume that a network consists of populations of neurons which share a similar external input and preferentially connect together with a homogeneous connection probability and strength. All connections originating from outside this circuit are considered to be feedforward input (see schematic Fig. **1A**, bottom). In biological circuits, it is difficult to determine what can constitute a single network since the brain exhibits a high degree of complexity and does not consist of well-separated circuits. Nonetheless, the analysis of cortical regions in which projection columns can be anatomically identified leads to a consistent order of magnitude for network sizes. Here we set the typical functional network to consist of 3000 E and 1000 I neurons. We present the corresponding citations below.

In the primary visual cortex of mice, it is especially difficult to define a local network size since the cortical map is unstructured, meaning that neurons which share the same receptive field do not co-localize [81]. However, this does not mean that the concept of homogeneous network cannot be applied to mouse V1, as neurons which share the same receptive field preferentially connect together [8]. Since local networks within mouse V1 cannot be defined based on spatial anatomy, we used other regions with anatomically-defined networks to define a reference point.

In mouse somatosensory cortex, distinct neuroanatomical structures known as barrels receive the sensory input from each whisker [4]. These structures are perfect candidates to define the typical scale of a homogeneous network. Their diameter ranges from 100 to 400 *µ*m, with a thickness in layer 4 of 100 *µ*m [4]. Using the neuronal density observed in [5], the number of neurons in these structures ranges from 140 to 2200. In the rat barrel cortex, the number of neurons in each layer of multiple projection columns has specifically been counted [3], and is on the order of *N* = 4000 in L4 and *N* = 6000 in L2/3. In primates, the primary visual cortex of macaques has been studied extensively. Unlike rodents, the cortical organization of macaque V1 shows a columnar structure, both for eye dominance and orientation preference [82]. Within a range of orientation preference of 10^*°*^, such columns are slab-shaped, with a size of 30 *µ*m by 0.5 to 1mm [83]. With the thickness of L2 and L3 being respectively 225 *µ*m and 310 *µ*m [84], and a cell density of 1.3 ×10^5^ neurons*/*mm^3^ [85]. We can deduce that the number of neurons in L2/3 in such columns is in the range of 1000 to 2000.

In summary, it appears that across species and cortical regions, we can define functional networks with sizes ranging from hundreds to a few thousands of neurons. In this work, we choose a network size of 4000 because it corresponds to the value reported in [3], which is the only study where the neurons in a cortical column were directly counted. In this context, we can use anatomical analyses of neurons in the same brain region to determine the fraction of excitatory and inhibitory neurons in a network [7, 86], which leads to a E/I ratio of 3.5 in layer 2/3. This corresponds to 3110 E neurons and 890 I neurons, which we round to 3000 and 1000 respectively.

Finally, we verify that the network size we assume (i.e. 4000 neurons) is plausible for mouse V1, since we could not use a spatial anatomy feature to define a network. We use the reported probability profile (*p*) for E→E connections as a function of distance for mouse V1 L2/3, Fig. 4B in [54] (near 20% for nearby pairs, going down to 0% for neuron pairs 150*µ*m apart). Following this observation, the number of excitatory synapses to an excitatory neuron can be obtained through a 3D spatial integration of this connection probability profile: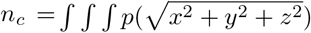, where *η* is the density of excitatory neurons. Over an infinite 3D space, we obtain the total number of E → E connections: *n*_*c*_ = 8.7 ×10^5^*η*. Using a neuronal density of 1.64 ×10^*−* 4^ neurons*/µm*^3^ [87], and assuming a E/I ratio of 3.5, we obtain *n*_*c*_ ≈ 111 connections. The result of this rough calculation is in the same order of magnitude as the 195 connections we obtain with 4000 neurons and a probability of connection of 6.5% (as described in [6]), which suggests that this network size is a valid approximation for mouse V1 as well. It should be noted that the network we define does not constitute a single block of cortex due to the salt-and-pepper organization of this brain region, but consists instead of distant neurons which receive the same external input and are homogeneously connected within the network.

## Appendix B: Derivation of parameter conditions for specific computational regimes

### 1. Balanced state framework

Here, we provide for completeness the solutions of the balanced state framework derived previously [16] which we used as a reference to study the convergence to the balanced state. In this framework, the mean input to each population vanishes as the number of neurons in the network increases, *µ*_*E*_ ≈ *µ*_*I*_ ≈ 0 and the equations for *µ*_*E*_ and *µ*_*I*_ in Eq. **12** can be simplified to

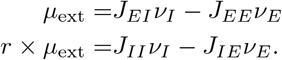

The solution of the balanced state equations reads

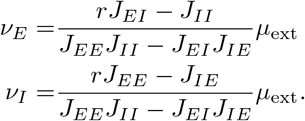

The balanced state solution is only valid if both *ν*_*E*_ and *ν*_*I*_ are positive for positive input, which corresponds to the following condition on connectivity

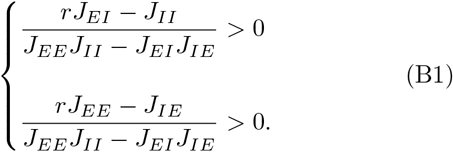

#### Stability of the balanced state

Within the balanced state framework, we assume that the change in firing rate of each population is a function of the excess input it receives.

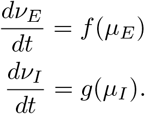

The firing rate of a population is at the steady state when its total input is balanced (*f* (0) = 0 or *g*(0) = 0). By linearising around a steady state, we get

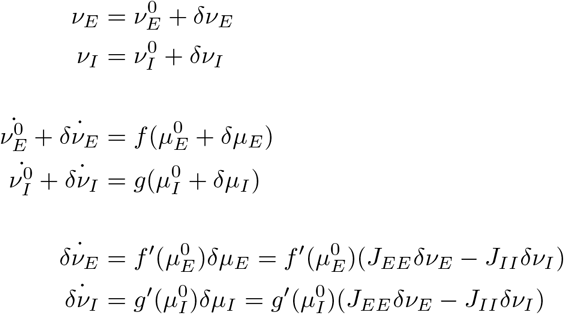

This can be rewritten as

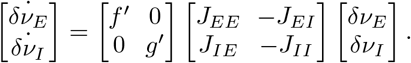

Where *f*′ and *g* ′ are positive (excess input drives the firing rate up), the state 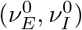 is stable if the two eigenvalues of the Jacobian matrix

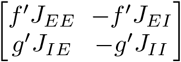

have negative real parts.

The eigenvalues *λ*_1_ and *λ*_2_ are roots of the polynomial

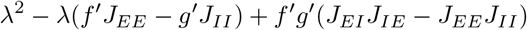

or

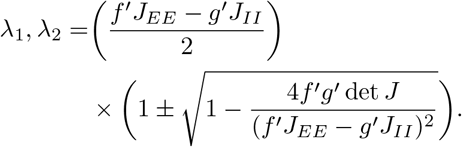

The steady state is stable ⇔

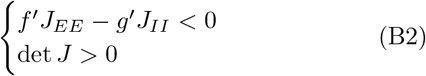

The first condition requires that the response of the inhibitory population (*g*′) is fast and strong enough to prevent a runaway excitatory feedback loop. We do not use this condition here because it depends on the dynamic properties of the network (*f* and *g* functions), which are beyond the scope of this work. However, the second condition constrains the connectivity matrix such that *J*_*EI*_*J*_*IE*_ − *J*_*EE*_*J*_*II*_ *>* 0 [24].

The condition on the existence of a non-negative balanced state (Eq. **B1**) can be combined with the stability condition on connectivity (Eq. **B2**) to delineate the parameter range where a balanced state limit exists and is stable [16, 24]:

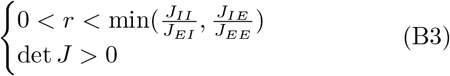

We use Eq. **B3** to visualize the parameter range where a balanced state exists in Fig. **5** and Fig. **S1**.

### 2. Condition on supersaturation

Supersaturation is characterized by a decrease in the excitatory firing rate with increasing external input: 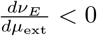.

Linearizing the system around a fixed point, leads to the relation between input and firing rate at the steady state [47]:

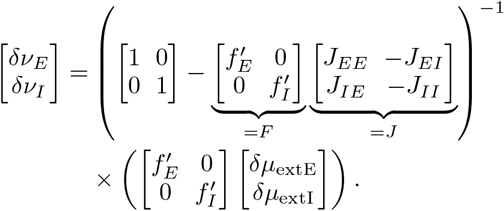

Where *δν* and *δµ*_ext_ are the firing rates and external inputs linearized around a steady state. The functions *f*_*E*_ and *f*_*I*_ are the input-firing rate transfer functions of the two populations. The functions 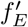 and 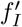 are the derivatives with respect to input, calculated at the fixed point. or The effect of a change of external input yields:

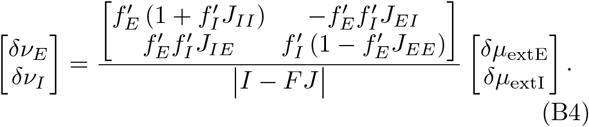

In particular, the effect of external input on the excitatory firing rate yields

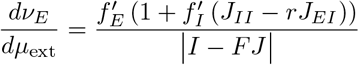

As shown in [47], |*I* − *FJ*| must be positive for the fixed point to be stable. Furthermore, 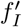 and 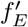 sumed to be positive, meaning that the F-I curves are monotonically increasing. This leads to the following condition for supersaturation:

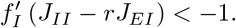

Which is only possible if 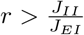. In the SSN, the transfer function is a power law (Eq. **1**). Its derivative is then 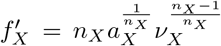. This leads to the condition that supersaturation occurs when the fixed point inhibitory firing rate is sufficiently high [15]:

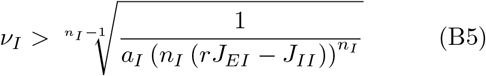

For completeness, it is worth mentioning that there exists one particular edge-case where 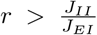 cannot not lead to 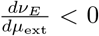. This occurs when the only stable state of the system is such that *ν*_*E*_ = 0 regardless of *µ*_ext_. This is discussed in the section *Effect of J*_*EE*_ *on the occurence of ISN* and requires *rb*_*E*_ − *b*_*I*_ *>* 0. Since these cases correspond to situations where recurrent inhibition is strong enough to prevent any excitatory activity, we consider them to be supersaturating.

#### Modulation of E firing rate peak in supersaturating activity regime

Here, we explain how we modified the height of the E firing rate peak in Fig. **2B**. We begin by analyzing how the value of maximal E firing rate depends on SSN parameters for supersaturating networks. A characterizing property of the maximal E firing rate *ν*_*E*_ is that it satis-fies 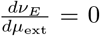. As shown previously [15], this occurs for 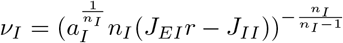, where *J*_*EI*_*r* – *J*_*II*_ must be positive. Since *ν*_*I*_ and *ν*_*E*_ are both positive, we can remove the (·)_+_ operator in the SSN equations and apply the inverse power law exponents to both sides to express *µ*_ext_ Eq. **3**

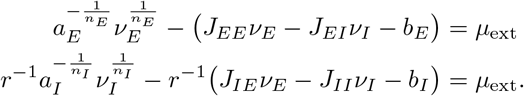

We combine two above equations to obtain

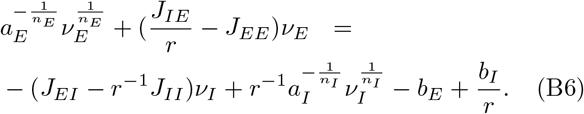

Substituting 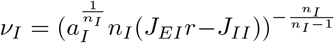 in the above equation, we obtain

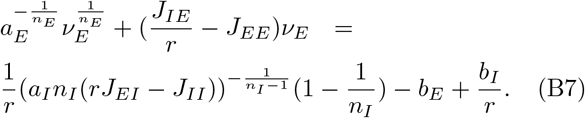

The solution of Eq. **B7** corresponds to the maximal E firing rate in the supersaturating activity regime. To increase the E firing rate peak, we modified *r, J*_*IE*_, and *J*_*EI*_. Specifically, we decreased *r* and modified *J*_*IE*_ and *J*_*EI*_ such that the terms *J*_*IE*_*/r* and *J*_*EI*_*r* remained constant. If *J*_*IE*_*/r* − *J*_*EE*_ is positive, the left-hand side of the equation is a monotonically increasing function of *ν*_*E*_. As 1*/r* increases, the right side of Eq. **B7** moves upward and the corresponding *ν*_*E*_ on the left side must increase as well. In this way the unique solution *ν*_*E*_ of Eq. **B7** - the maximal E firing rate - increases as 1*/r* increases. The method demonstrated here assumes that the left hand side of Eq. **B7** is an increasing function of *ν*_*E*_. This is the case if *J*_*IE*_*/r* − *J*_*EE*_ is positive (as in Fig. **2**), and the peak of the excitatory activity can be increased infinitely. On the other hand, if *J*_*IE*_*/r* − *J*_*EE*_ is negative, the peak of supersaturation is bounded. In particular, for supersaturating networks (*J*_*EI*_*/r*− *J*_*II*_) for which det *J* is negative, *J*_*IE*_*/r* − *J*_*EE*_ is negative. The peak of supersaturation is therefore bounded, and decreasing *r* can lead to an unstable situation where no steady state exist (as illustrated in Fig. **5B**).

### 3. Condition on the paradoxical effect

The paradoxical effect [25, 47, 48] is characterized by a decrease of the I firing rate, as the external input to the I population is increased: 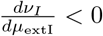. Here again, the effect of a change in external input is given by Eq. **B4** where | *I* − *FJ* | and 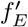 must be positive. In this case, 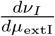 can only be negative if

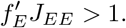

This condition is equivalent to the condition for the instability of the excitatory subnetwork [47]. The paradoxical effect is therefore a feature of the inhibition stabilized network (ISN) as it occurs when the activity of the E population is only made stable thanks to the suppression from the I population. Using the SSN transfer function ^*−*^ (Eq. **1**), the condition on the paradoxical effect leads to Eq. **7**

#### Effect of J_EE_ on the occurence of ISN

Since the network is inhibition-stabilized for fixed points which satisfy Eq. **7**, it seems that any network can be in the ISN state, granted that *J*_*EE*_ is sufficiently strong. Here we ask whether there is any counterexample to this. Are there networks which cannot enter ISN, no matter how high *J*_*EE*_ is?

From Eq. **B6**, we impose that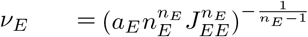, which corresponds to the onset of inhibition-stabilization to get

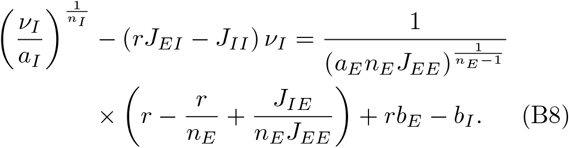

This equation can be seen as the crossing of two functions where the left hand side *f* is a function of the I activity *ν*_*I*_ and the right-hand side *g* is independent of *ν*_*I*_ but varies with *J*_*EE*_.

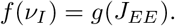

If there is a value of *ν*_*I*_ which satisfies this equation, the network has a fixed point such that *ν*_*E*_ is at the onset of ISN. We note from Eq. **B6** that

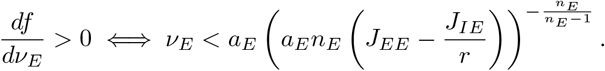

Since the value of *ν*_*E*_ at which we operate in Eq. **B8** is smaller than this (for *J*_*IE*_ ≠ 0), the network cannot be in the ISN if *f* and *g* do not intersect and *f* remains below *g*.

The *f* function starts at 0 for *ν*_*I*_ = 0 and is either monotonously increasing (if *J*_*II*_ *> rJ*_*EI*_) and tends to ∞, or it reaches a maximum and then tends to − ∞ (if *rJ*_*EI*_ *> J*_*II*_). The second case corresponds to supersaturation, where high values of *ν*_*I*_ can be sustained with low recurrent excitation. Whatever the parameters, *f* first rises and always has at least some positive values.

The *g* function can be represented as a polynomial

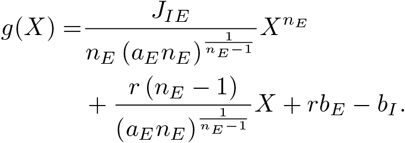

Where 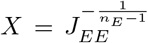. Since the coefficients in front of 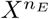 and *X* are both positive and X is necessarily positive (because *J*_*EE*_ is positive), *g* can take all values larger than *rb*_*E*_ − *b*_*I*_. This means that, with *J*_*EE*_ as a free parameter, equation Eq. **B8** will have a solution if there is any *ν*_*I*_ such that *f* (*ν*_*I*_) *> rb*_*E*_ − *b*_*I*_.

For values of *b*_*E*_ and *b*_*I*_ such that *rb*_*E*_ − *b*_*I*_ *<* 0, the network can always enter an inhibition-stabilized state by tuning *J*_*EE*_. Interestingly, this is always the case with the values of *b*_*E*_ and *b*_*I*_ we obtained from fitting the F-I curve (See Table **III**) because experimentally reported neuronal parameters [6] are such that *τ*_*E*_ *> τ*_*I*_. Similarly, in networks which do not satisfy the supersaturation condition (*J*_*II*_ *> rJ*_*EI*_), *g* always crosses *f*.

On the other hand, in networks for which *f* has a maximum (*J*_*II*_ *> rJ*_*IE*_), if *rb*_*E*_ − *b*_*I*_ is higher than this maximum, the functions *f* and *g* never cross regardless of the value of *J*_*EE*_. In this scenario, the network can never reach a steady state where it is inhibition-1 stabilized. These cases correspond to situations where the E activity is always suppressed and there is no stable steady state with *ν*_*E*_ *>* 0. If *J*_*EE*_ is large, another unstable steady state will exist (for det *J <* 0, see section *Parity of solutions* below), and if the system is perturbed enough to reach it the E activity will enter an unlimited feedback loop which leads to always increasing activity. These systems cannot be in the ISN regardless of *J*_*EE*_ because the E activity is either entirely silent or cannot be stabilized by inhibition.

### 4. Condition on multiplicity of solutions

As shown in [28], the two-dimensional SSN equation (Eq. **3**) can be rewritten as a single characteristic function ℱ, where the steady states of the system correspond to zeros of ℱ.

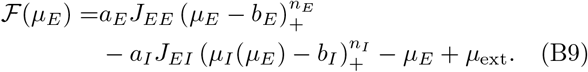

Where *µ*_*I*_ is a function of 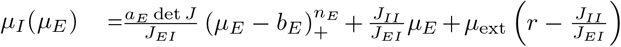. The first term of ℱ is zero if the E population is silenced (*ν*_*E*_ = 0), and the second one if the I population is silenced (*ν*_*I*_ = 0). For any value *µ*_*E*_ which satisfies ℱ (*µ*_*E*_) = 0, the corresponding excitatory firing rate is given by the power law F-I function Eq. **1**.

The number of zero crossings of the function ℱ corresponds to the number of fixed points of the system. Since ℱ is a continuous function, its number of zero crossings only changes when two solutions merge into one or when one solution splits into two. This corresponds to the situation when extrema of ℱ fall on zero.

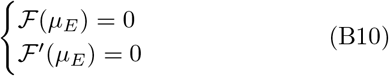

Where ℱ ′ denotes the derivative of ℱ with respect to *µ*_*E*_. This condition corresponds to changes in the number of solutions. The parameters comprised between these boundaries have the same number of solutions. This approach can be used to delimit the range of bistability or absence of solutions (as shown in Fig. **5**). Within such a region, the number of network states is obtained by determining the number of zero crossings of ℱ.

#### Parity of solutions

The number of zero crossings of the ℱ function can be studied through its limits. Assuming that the F-I curves are supralinear(*n*_*E*_ *>* 1 and *n*_*I*_ *>* 1), we get:

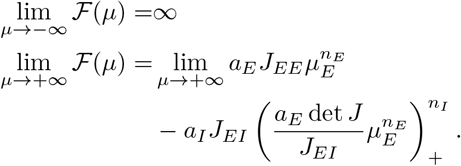

If det *J >* 0, the second limit tends to − ∞. Therefore, the function has at least one zero and the ℱ function for positive determinants has have an odd number of solutions (Mean-value theorem). On the other hand, if the determinant is negative, the second limit tends to − ∞. In that case, there is no guarantee that the system has a fixed solution and the number of solutions is even. Multiple roots (Eq. **B10**) are counted separately in this calculation.

## Appendix C: Noise contribution of exponential synapses

In this section we expand the mean-field approximation of the recurrent noise to the case of exponential synapses. In the case of the delta synapse (Eq. **10**), by assuming a Poisson spike train, the standard deviation is given by *σ*_*XY*_ = *j*_*XY*_ *J*_*XY*_ *ν*_*XY*_ (Eq. **12**). For exponential synapses however, since the current is distributed in time Eq. **11**, the noise is lower than this. The exponential synapse corresponds to a shot noise process, and its covariance is given by [88, 89]:

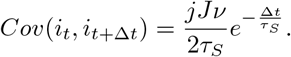

Over a period Δ*t*, the variance of the received current is

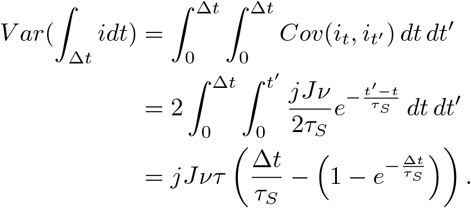

The Φ equation is derived from the analysis of a Ornstein Uhlenbeck process with white noise [39], so that the noise *σ* considered in Φ is the rate of increase of the variance. For exponential synapses, we then approximate a white noise Ornstein Uhlenbeck process by using the rate of increase of variance:

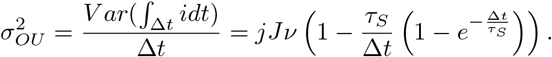

Since the current at different time steps is correlated, the variance of the overall cumulative current over a time period Δ*t* is not a linear function of Δ*t*. This contrasts with white noise processes where this rate is a constant.

For Δ*t* ≪ *τ*_*S*_, *σ*_*OU*_ tends to zero whereas for Δ*t* ≫ *τ*_*S*_, *σ*_*OU*_ approaches the Poisson limit *jJν*. In the context of the mean field analysis of the network, we set the period Δ*t* over which *σ*_*OU*_ is considered to be the period between two successive postsynaptic spikes. Therefore, we use the expected value of *σ*_*OU*_ over the distribution of ISI. For Poisson processes, the ISI follows an exponential distribution.

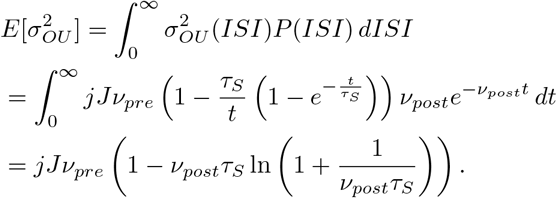

All in all, the noise originating from the recurrent exponential synapses can be quantified and the Ricciardi self-consistency solution Eq. **12** can be adapted to take this effect into account:

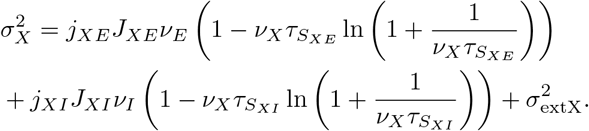

However, it should be noted that this approach can only provide an approximation of the firing of LIF neurons since the Φ function is only exact for uncorrelated white noise input.

## Appendix D: Supplemental figures

**FIG. S1.**
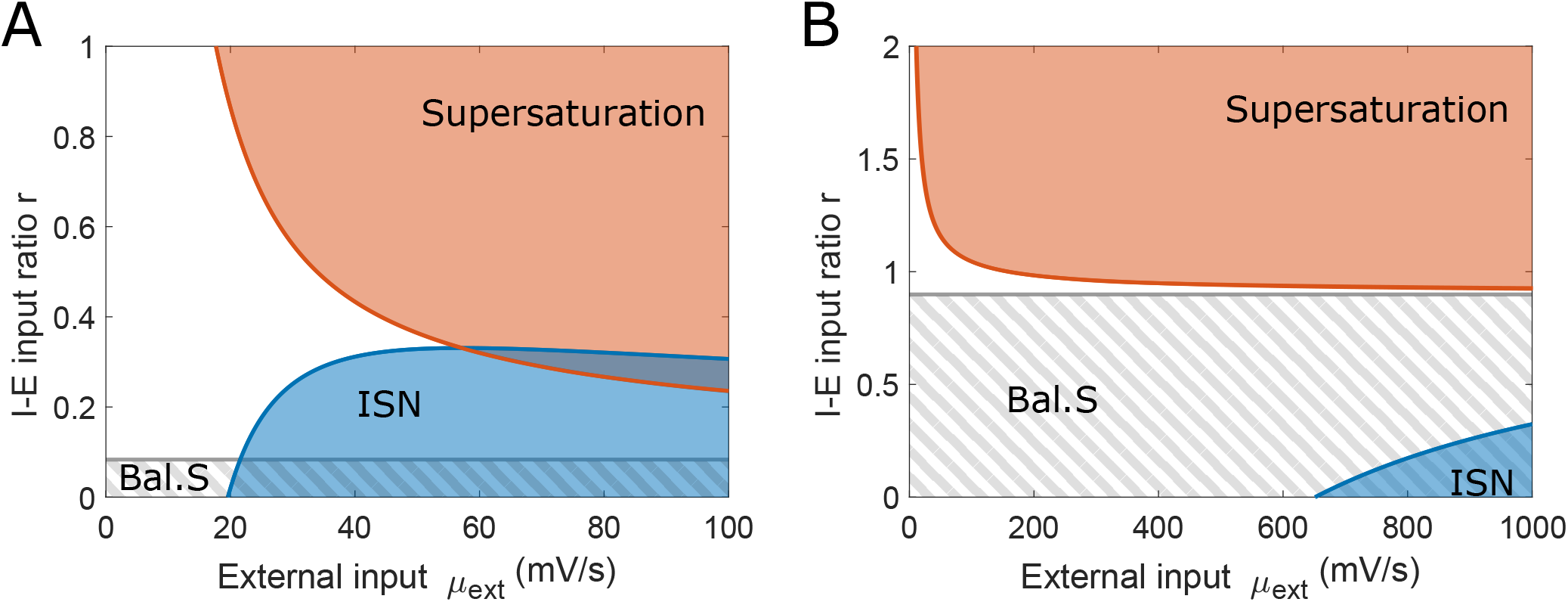
Additional map of computational regimes. These maps are equivalent to the maps shown in Fig. **5**, and are generated for the connectivity of the supersaturating network shown in Fig. **2** and the mouse V1 network shown in Fig. **3A** (A) The map shows many similarities to the map shown in Fig. **5A**. The balanced state is only defined for low *r* values across external input values. Te network can be inhibition stabilized for large input and low *r*, whereas supersaturation occurs for large input and high *r*. The supersaturation and ISN regions overlap. However, unlike Fig. **5A**, this network does not have a bistable regime in the range of inputs presented here.(B) Compared with the phase space in panel A, the ISN state (blue area) appears more difficult to achieve for this network as it requires much higher external input to reach. We show in Fig. **3B** that increasing *J*_*EE*_ makes the ISN accessible for external inputs *µ*_ext_ lower than 100 mV/s.

**FIG. S2.**
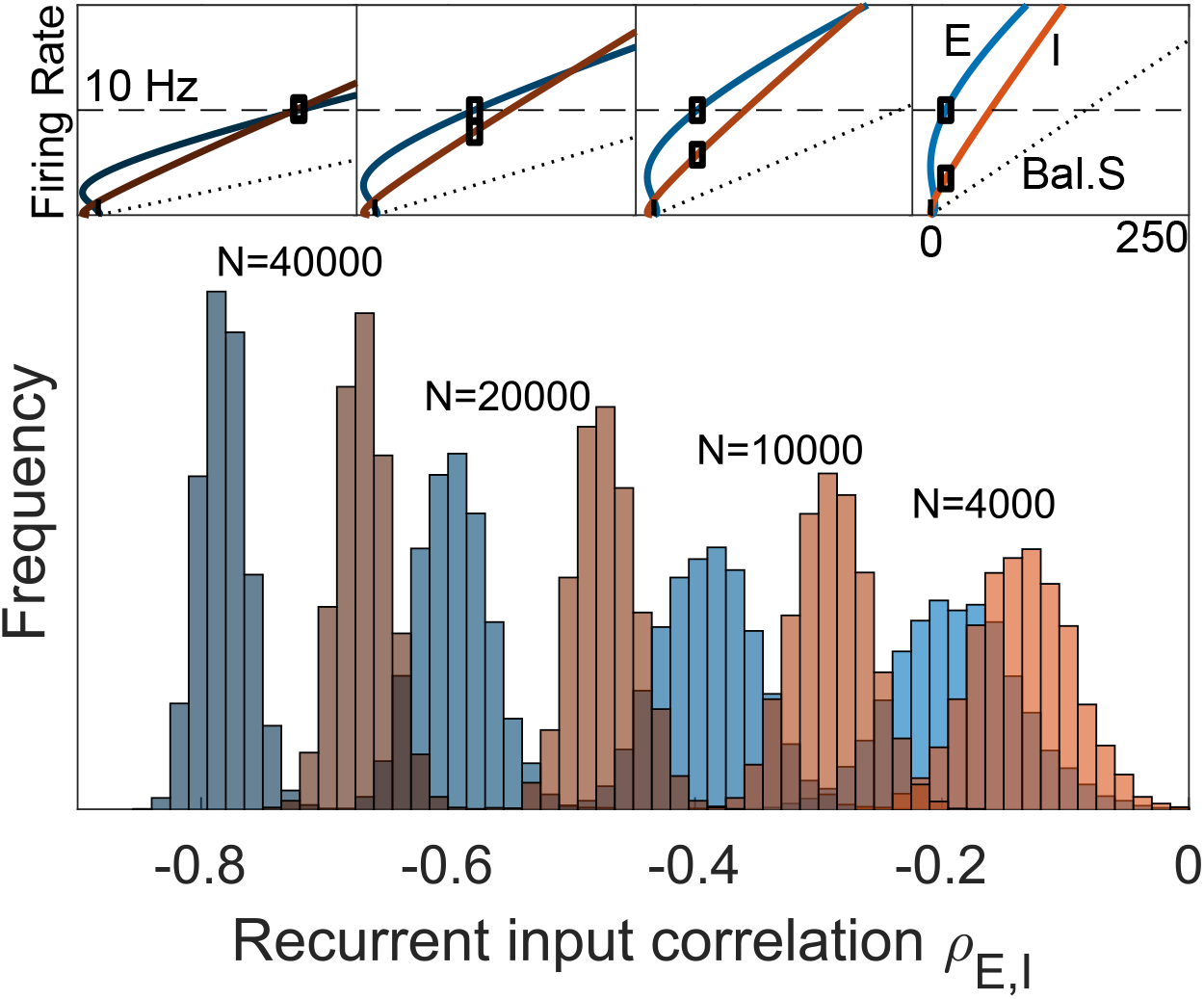
Effect of network size on E/I input balance. As the size of networks increases, the correlation between incoming E and I currents (*ρ*_*E,I*_) becomes stronger. This is measured in E (blue) and I (red) neurons for the same bistability case as Fig. **4** and Fig. **6C**, at the point at which the excitatory firing rate reaches 10Hz (see inset above, for each network size). This shows that the E-I balance gets tighter as the network size increases even though the firing rates are far from the balanced state limit (dashed line, inset) and the network remains non-linear.

